# Extraction of bouton-like structures from neuropil calcium imaging data

**DOI:** 10.1101/2021.05.28.445372

**Authors:** Kazushi Fukumasu, Akinao Nose, Hiroshi Kohsaka

## Abstract

The neuropil, the plexus of axons and dendrites, plays a critical role in operating the circuit processing of the nervous system. Revealing the spatiotemporal activity pattern within the neuropil would clarify how the information flows throughout the nervous system. However, calcium imaging to examine the circuit dynamics has mainly focused on the soma population due to their discrete distribution. The development of a methodology to analyze the calcium imaging data of a densely packed neuropil would provide us with new insights into the circuit dynamics. Here, we propose a new method to decompose calcium imaging data of the neuropil into populations of bouton-like synaptic structures with a standard desktop computer. To extract bouton-like structures from calcium imaging data, we introduced a new type of modularity, a widely used quality measure in graph theory, and optimized the clustering configuration by a simulated annealing algorithm, which is established in statistical physics. To assess this method’s performance, we conducted calcium imaging of the neuropil of *Drosophila* larvae. Based on the obtained data, we established artificial neuropil imaging datasets. We applied the decomposition procedure to the artificial and experimental calcium imaging data and extracted individual bouton-like structures successfully. Based on the extracted spatiotemporal data, we analyzed the network structure of the central nervous system of fly larvae and found it was scale-free. These results demonstrate that neuropil calcium imaging and its decomposition could provide new insight into our understanding of neural processing.

## 1. Introduction

Calcium imaging of the cell bodies of neurons has provided rich information on the functional dynamics in neural circuits. Functional imaging of neuron populations has revealed the existence of maps in several regions of the central nervous system (CNS), including the visual cortex, auditory cortex, hippocampus, amygdala, and spinal cord (Gauthier & Tank, 2018; Gründemann et al., 2019; Hackett et al., 2011; Tinnermann et al., 2021; Zhuang et al., 2017). The configuration of cell bodies is one of the critical factors for neural circuit computation.

While the calcium imaging of somas has been widely used, the cell body is not the sole component of a neuron. For synapse transduction, axons and dendrites are major structures forming the brain’s neuropil. Considering the physiological roles in the neuropil, pan-neuronal calcium imaging of the neuropil would bring us several new types of information on the circuit dynamics (London & Häusser, 2005; Poirazi & Papoutsi, 2020). First, the calcium signal in the neuropil directly reflects the synaptic transmission, which is critical to understanding spatiotemporal dynamics in the nervous system. Second, the activation of one neuron could generate heterogeneous activity among its multiple presynaptic sites (Bilz et al., 2020). This heterogeneity can be detected by neuropil imaging but not by somatic imaging. Third, since the neural wiring diagram is embedded in the neuropil, the activity dynamics in the neuropil can represent the flow of information in the neural circuit (Kohsaka et al., 2019; Kreiner et al., 1987; Lee et al., 2017; Moyle et al., 2021).

Whereas neuropil imaging would be informative in studying circuit dynamism, the data analysis of calcium imaging in the neuropil has been challenging. Because synapses and neurites are densely packed in the nervous system, it is hard to identify synapses and extract their activity from pan-neuronal imaging data. A variety of algorithms to decompose calcium imaging data into cellular activities have been developed based on several mathematical approaches, including principal component analysis, independent component analysis, similarity analysis, singular value decomposition, non-negative matrix factorization, and graph theory (Maruyama et al., 2014; Mölter et al., 2018; Mukamel et al., 2009; Pnevmatikakis et al., 2016; Shibue & Komaki, 2020). However, automatic approaches to decompose the neuropil remain to be developed. Especially most of the decomposition algorithms require that activities of neuronal populations are sufficiently sporadic and independent so that signal traces of proximate cell bodies can be clearly differentiated. However, synapse population activity patterns in the neuropil are not always sporadic but spatiotemporally coordinated in some cases, such as global bursts and wave-like propagations (Churchland et al., 2012; Lemon et al., 2015). Therefore, the existing algorithms might not be suitable for processing calcium imaging data with spatially coupled activities.

In this study, we propose a new methodology for analyzing neuropil calcium imaging data and an unsupervised algorithm to extract pre-synaptic bouton-like structures and their activity traces. We apply an iteration method to adaptively infer the baseline of the signals in spatial dimensions, which enables the elimination of coarse signals from out of the focal plane and the extraction of well-focused signals. To label each bouton-like component in the pre-processed data, we introduce a new type of modularity measure (Humphries, 2011; Newman & Girvan, 2004) and optimize the clustering configuration by a simulated annealing algorithm (Kirkpatrick et al., 1983; Metropolis et al., 1953), which is established in statistical physics. We apply this procedure to artificial and biological calcium imaging data and successfully extract population activity in the neuropil for motor control showing wave-like activity patterns.

## 2. Methods

In this study, we developed a novel methodology for analyzing neuropil calcium imaging data requiring low computational resources. We realized this by taking advantage of the observation that each bouton had a compact, round shape. By omitting unnecessary computation between distant pixels, we built an effective decomposing algorithm with high sensitivity.

We used python3.7 combined with C++ and python libraries, NumPy, SciPy, and OpenCV on a standard desktop computer (maximum speed: 3.60 GHz, 4-core CPU, 8 GB RAM) for the calculation.

### 2.1. Patch-wise modularity-based pixel clustering

To extract bouton-like structures, we defined a patch-wise similarity matrix based on inter-pixel signal similarity, obtained a patch-wise modularity matrix, and solved the optimization problems using the Gibbs sampling.

#### 2.1.1. Similarity matrices

First, we applied pixel clustering based on comparing the temporal profile of pixels’ signal intensity. This procedure was devised with the idea that the profile of the calcium signal from the points within the same bouton should be similar. Because boutons were densely packed in the neuropil, it was difficult for the computer to recognize them solely by their morphological features. Accordingly, we utilized their temporal information for clustering. However, it was still difficult to compare the pixels in pairs because the number of the pixel pairs was too large (about 10^12^ pairs) for standard desktop computers. Instead of analyzing every pair, we compared pixel pairs whose distance was within a patch radius r (section 2.1.2). The patch radius was set to 3-6 pixels. By so doing, the number of the comparisons could be small enough to handle with standard desktop computers, and bouton-sized regions (the diameter of about five pixels) were extracted (section 2.1.3).

To define the similarity for clustering, we referred to experimental observation. The calcium signals of pixels within each single bouton changed synchronously and exhibited no apparent time lag among pixels (Fig. C.1.A-C). Therefore, as a temporal similarity between two pixels *μ* and *ν* (*Ã_μν_*), we adopted the inner product between the normalized data vectors,

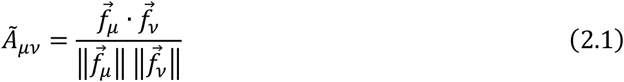

where 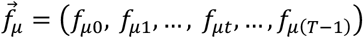 (*μ*: pixel, *t*: frame) was a vector of signal intensity, and ||·|| indicated the Euclidean norm. In the following analysis, we regarded the normalized inner products as weights in a graph. Although most (>99.99%) of the values were non-negative in our calcium imaging datasets, a few values were negative. Accordingly, if the value was negative, we replaced it with zero. In the end, we defined the similarity matrix *A_μν_* as follows:

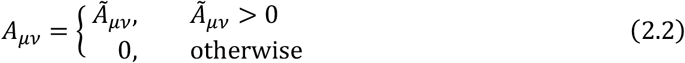

#### 2.1.2. Calculation of patch-wise modularity matrices

By using the similarity matrix, we calculated the modularity matrix *B_μν_* that described a set of clusters of components based on their similarity. Conventionally, the modularity matrix was defined as the following (Humphries, 2011; Newman, 2006):

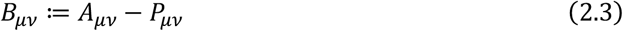

where *P_μν_* was the expected value of similarity for weight-randomized networks keeping the same degree distribution with *A_μν_ (∑_ν_A_μν_ = ∑_ν_P_μν_*). In the randomization step, the similarity value of an edge was supposed to be determined by the node degree at the two ends of the edge. Accordingly, *P_μν_* was described as the product of the selection probabilities of two nodes and the total edge weight. The selection probability of node *μ* was *d_μ_*/*m* and the total edge weight was *m*, where *d_μ_*:= ∑_*ν*_*A*_*μν*_, *m*:=∑_*μν*_*A*_*μν*_ = ∑_*μ*_*d*_*μ*_. Therefore, *P_μν_* = *d_μ_/m*· *d_v_/m·m* = *d_μ_d_v_/m* was derived. From the definition of the modularity matrix (Eq. (2.3)), *B_μν_* indicated the extent of similarity between *μ* and *ν* compared with the expectation. With a community configuration *S* = {*s_μ_*| *μ*: pixel, *s_μ_*: community id of pixel *μ*}, the quality function termed modularity was defined as follows:

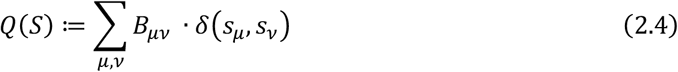

where *δ* was the Kronecker delta function *(δ(x,y) = 1* if *x = y,* and 0 otherwise). Because *B_μν_* was added to the modularity *Q* if *μ* and *v* were in the same community (therefore, the modularity was upregulated or downregulated according to whether *B_μν_* was positive or negative, respectively), *B_μν_* could be assumed as the interaction intensity between components (attraction if positive and repulsion if negative). Therefore, the optimized community configuration that maximized the modularity divided components into communities most clearly, so that the components were more similar within them and less similar between them.

However, the conventional modularity matrix could not be used for our pixel clustering. First, it was practically impossible to calculate all components of *A_μν_* because the number of pixel combinations was too large to handle. The method needed to hold connectivity strength between every combination of nodes, which in our case involved data for more than two million pixels, a task that was impossible to carry out on a standard computer. Second, modularity-based community detection had problems with the “resolution limit” (Fortunato & Barthélemy, 2007). Since the expected number of links in a random network decreased in a large network, the elements in the modularity matrices tended to be large, and clusters would be merged in a large network. This effect could decrease the spatial resolution of the clustering and make bouton-like structures merge.

To engineer a modularity-based pixel clustering method, we defined “patch-wise modularity” by modifying the conventional definition (for the detailed definition, see Appendix A). The concept of patch-wise modularity was based on the idea that only spatially close pixels should be compared to recognize bouton-sized regions. At first, we introduced a weight matrix to restrict the similarity calculation to neighborhoods as pixel-centered circle patches:

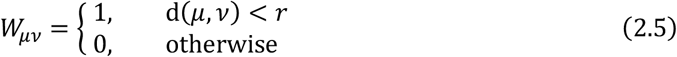

where d(*μ,ν*) was the Euclidean distance between pixels *μ* and *ν*, while *r* was the radius of patches that determined the scale of output pixel clusters. The patch radius *r* was chosen from 3.0-5.0 pixels, a typical diameter of a single bouton in our calcium imaging data. Instead of *A_μν_*, we used *Â_μν_* = *W_μν_:= A_μν_* as the similarity matrix. Since the number of the non-zero components in *Â_μν_* was reduced drastically compared with *A_μν_*, the calculation and memory costs became to be manageable size (about 1/3,000 times smaller than the original one). Then, we calculated 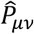, the expected similarity of networks with the same degree distribution 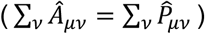. Unlike the conventional 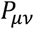, we additionally imposed a localization condition, 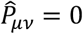 when *d*(*μ,ν*) ≥ *r*, which was necessary to ensure the modularity within each local region. However, the conventional definition 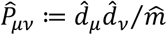, where 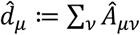 and 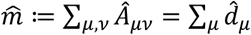, could not generally fulfil the localization condition. Therefore, we expanded the conventional definition to meet the localization condition and obtained 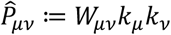, where *k_μ_* was a vector chosen to satisfy 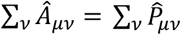. For given *W_μν_* and *Â_μν_,* there always existed one and only one *k_μ_* (Theorem 1, Appendix B). Because *k_μ_* could not be described explicitly, we approximated 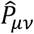 by an iteratively updated matrix according to the following recurrence formula:

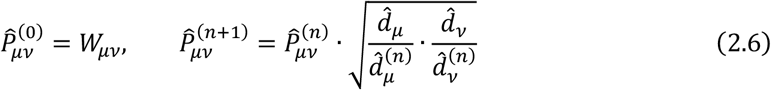

where 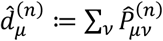. It was proved that the limit 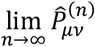 gave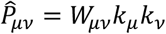 where 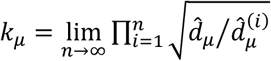 satisfying 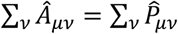 (Theorem 2, Appendix B). Therefore, we put 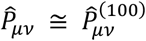 which had sufficiently converged to a specific matrix. Then, the patch-wise modularity matrix was calculated as follows:

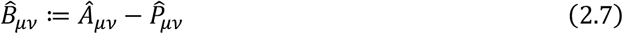

For the following optimization step, 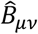 was divided by the standard deviation of 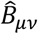 for pairs *μ* and *ν* where *W_μν_* = 1 to normalize the convergence speed in the optimization process.

#### 2.1.3. Optimization of the patch-wise modularity

Patch-wise modularity 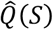 was defined from 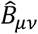 as an extension of Eq. (2.4), and 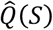 was maximized by optimizing the configuration of pixel clusters *S* = {*s_μ_*}:

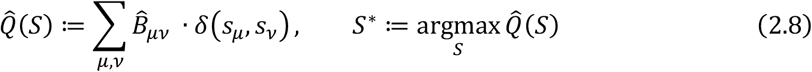

where 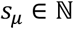 (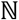 denoted natural numbers). The optimized configuration S* divided pixels into clusters based on the proximity and activity similarity.

To maximize the patch-wise modularity, we adopted simulated annealing (SA) method (Kirkpatrick et al., 1983) (Fig. 1.A), which was computationally efficient, especially when the interaction between elements (pixels in our case) was localized. In SA, the system was developed randomly according to the thermal state distribution with its energy (in our case, the energy 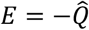) at a specific temperature. As the temperature gradually approached absolute zero, the system finally settled into the minimum energy state (then, 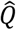 is maximized). The thermal state distribution was produced by iterative system update by the Gibbs sampling. The calculation cost of the Gibbs sampling was drastically reduced if the interaction among elements was localized.

**Fig. 1.**
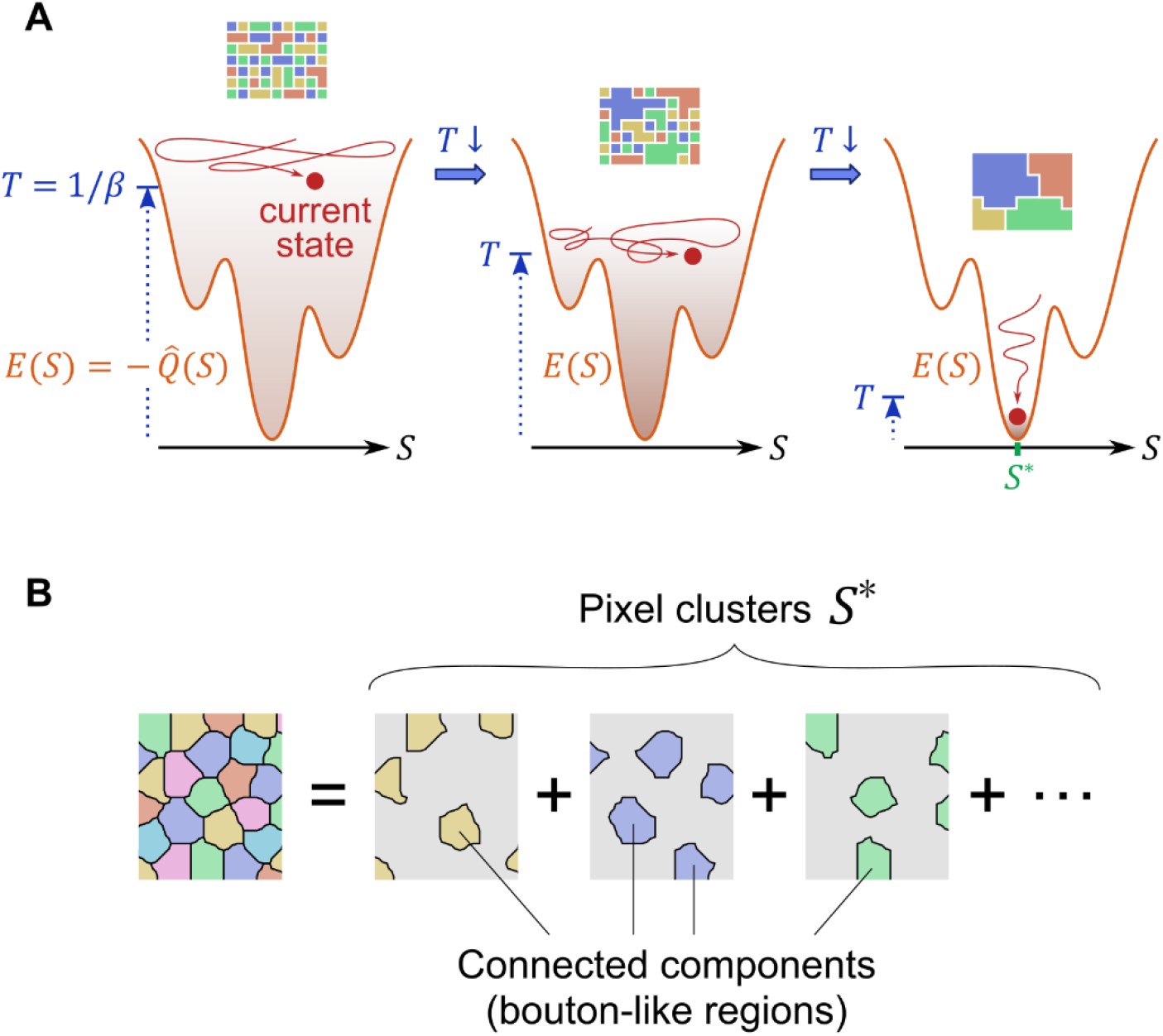
Optimization and extraction of bouton-like regions. (A) Schematic diagram explaining how simulated annealing works. The bottom three energy landscapes show that the energy is minimized, and the modularity 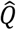 is maximized by searching the configuration space *S* as the temperature falls. The top three panels represent that the pixels in the image are clustered through the optimization process, as shown by enlarging domains of different colors. (B) The source image is decomposed into a set of pixel clusters, *S*\*. Each pixel cluster consists of a group of connected components, the diameter of which is around the threshold (patch radius r). Bouton-like regions are extracted as connected components in each of the pixel clusters in *S*\*.

For each iteration during the optimization processing with SA, the cluster identities of pixels *s_μ_* were updated one by one. At first, each pixel was randomly assigned with one of the labels whose number was the number of pixels within a patch of the radius r. This label number was sufficient to assign different clusters with distinct labels locally. Note that since the label number (about 100) was smaller than the total cluster number (about 5,000), each cluster could have multiple connected regions. Each single connected region would be isolated at the later step (see section 2.1.4.) Even when the total label number was set to larger numbers (200 and 400), the results were consistent with the case of 100 (Fig. C.1.D-F). The new *s_μ_* of each pixel was sampled according to the conditional probability distribution given by the Gibbs sampling, while identities of the other pixels were fixed:

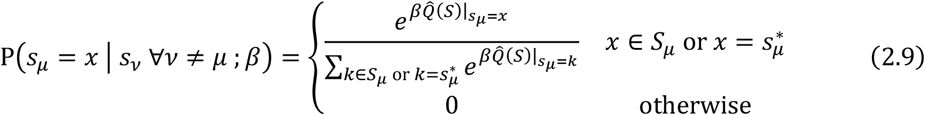

where *β* was the inverse temperature, *S_μ_* = {*s_ν_*|*d*(*μ,ν*) <*r, μ*≠*ν*} was cluster identities of pixels within the patch around the pixel *μ* except for *μ* itself, and 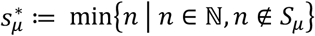. At each step, a candidate cluster label *s_μ_* for pixel *μ* was selected from labels around pixel *μ* (*S_μ_*) or a label not around it 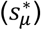. If the relabeling increased the pair-wise modularity 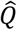, the candidate label was accepted, and the clustering configuration was updated with a high probability defined by Eq. (2.9). Because of the localized connectivity, most terms in the score 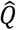 were independent of new *s_μ_* in Eq. (2.9). Accordingly, the formula of the probability was reduced to the following:

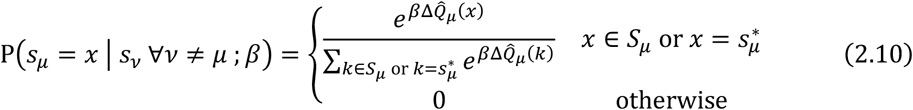

where 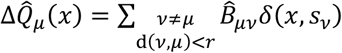. The number of terms in 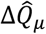 was highly smaller than that in 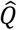, so the calculation cost for Eq. (2.10) was reduced from that of Eq. (2.9) by the ratio of 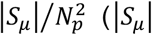: the number of elements in *S_μ_, N_p_*: the number of pixels). Within one iteration of SA, the identity of every pixel was updated according to the conditional probability distribution (Algorithm 1). The order of pixel update was randomly shuffled at every iteration (shuffle({1,2,…,*N_p_*})). Along with the iterations, the inverse temperature *β_i_* was raised. Because *B_μν_* was normalized by the standard deviation of its elements, 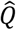 was almost converged when *β* reached 20 regardless of the data property, including pixel numbers. By comparing arithmetic or geometric progression of *β,* we found 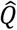 was maximized more successfully in the geometric case (Fig. C.1.G-I). Therefore, we raised *β* according to *β_i_* = 0.04 × 1.004^*i*-1^ (*i*. = 1,2,…), where *i.* is the iteration number. The configuration updates were terminated when *β_i_* reached *β*_max_ = 20 (*i*. = 1,557).

##### Algorithm 1. Simulated annealing method.

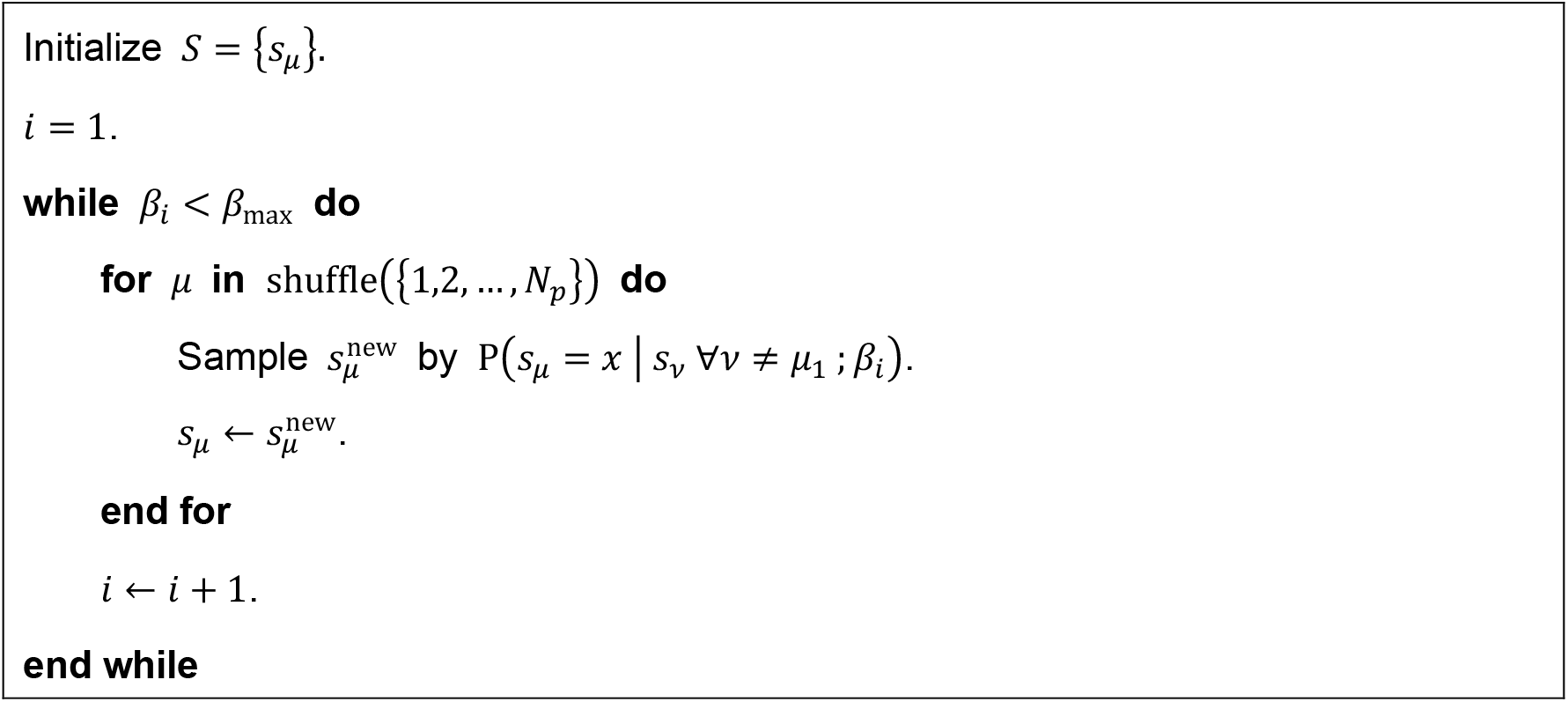

#### 2.1.4. Extraction of bouton-like regions

The optimized configuration *S*\* was used to define bouton-like regions in the neuropil imaging data. Each pixel cluster in *S*\* consisted of multiple connected regions (Fig. 1.B). However, since presynaptic boutons are a local and segregated structure, the multiple connected regions in a certain cluster could be regarded as distinct presynaptic regions. Therefore, *S*\* was further decomposed into spatially connected components {*R_n_*|*n* = 1,2,…,*N_C_*} (*N_c_*: the number of connected components), and the signals *f_μt_* in each *R_n_* were averaged 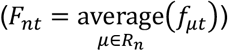.

#### 2.1.5. Selection of active bouton-like regions

Some of the bouton-like regions extracted from the calcium imaging data of the fly larval CNS were located outside of the ventral nerve cord and had almost no calcium signal. To filter out inactive regions from the data, we conducted threshold processing in terms of the S/N ratio of *F_nt_*. To evaluate the S/N ratio, the following two quantities were calculated:

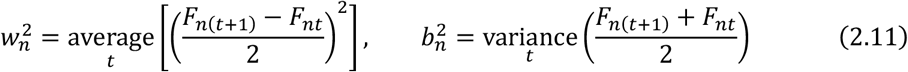

Here, *w_n_* was the average of the deviation of adjacent data points, and *b_n_* was the deviation of the average of adjacent data points. While *w_n_* measured the fluctuation between the consecutive frames, the fluctuation was averaged out in *b_n_*. Therefore, we regarded *w_n_* and *b_n_* as measures of noise and signal, and assumed the fraction *b_n_/w_n_* as an S/N ratio indicator. The larger *b_n_/w_n_* was, the better the S/N ratio was (white noise shows *b_n_/w_n_* = 1). Bouton-like structures were determined by choosing regions with *b_n_/w_n_* larger than 5.0 (Fig. C.2.A-C). The pixels in each extracted bouton-like region showed relatively similar calcium signal profiles (Fig. C.2.D). Furthermore, the size of the bouton-like structures was consistent with that of presynaptic boutons measured by immunostaining against presynaptic marker (Fig. C.2.E). To sum, the extracted regions would correspond to presynaptic structures in the neuropil.

### 2.2. Data acquisition and pre-processing

#### 2.2.1. Calcium imaging of the neuropil

Calcium imaging data were obtained from the CNS of third instar larvae in *Drosophila melanogaster* (Fig. 2.A). The genotype was *UAS-CD4::GCaMP6f; nSyb-Gal4*, which expresses a membrane-bound form calcium sensor (CD4::GCaMP6f) in all neurons *(nSyb-Gal4:* Bloomington #58763, *UAS-CD4::GCaMP6f:* (Kohsaka et al., 2019)).

**Fig. 2.**
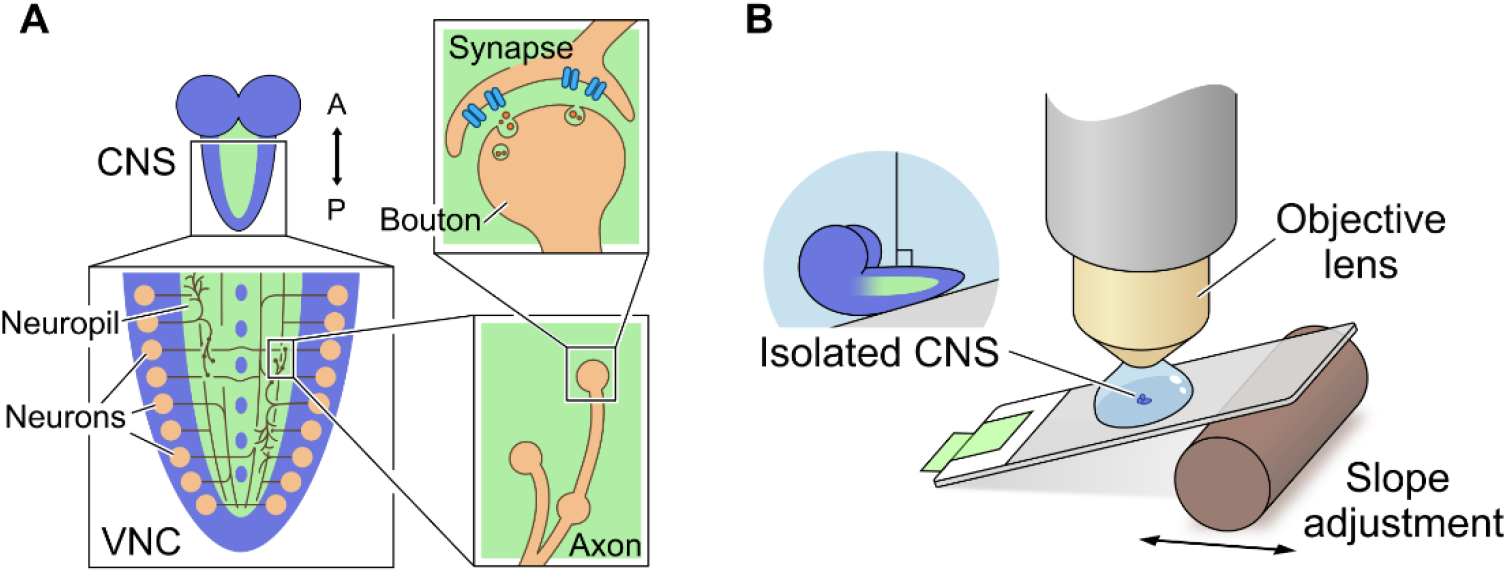
Calcium imaging of the VNC neuropil in an isolated CNS. (A) Schematics of the CNS, the VNC, the neuropil, and boutons in the *Drosophila* larvae. (B) Schematic diagram illustrating the setup for calcium imaging of the isolated CNS.

Calcium imaging of the CNS of fly larvae was carried out as described previously (Kohsaka et al., 2014, 2019) (Fig. 2.B). In brief, third-instar larvae of the *UAS-CD4::GCaMP6f; nSyb-Gal4* fly were dissected with microscissors, and the CNS was isolated. The isolated CNS was mounted on an adhesive slide glass (MAS coated slide glass S9215, Matsunami Glass, Japan) and soaked in an insect saline, TES buffer (TES 5 mM, NaCl 135 mM, KCl 5 mM, CaCl2 2 mM, MgCl2 4 mM, sucrose 36 mM; pH = 7.15). The fluorescence signal from the CNS was recorded by an EMCCD camera (iXon+ DU-897E-CS0-#BV, Andor, UK) (63x, 0.27 μm x 0.27 μm /px, 330 ms/frame, EM gain: 144, data depth: 14 bit) with attached spinning disk confocal unit (CSU21, YOKOGAWA, Japan). The CNS preparation was illuminated by a blue laser (power: 300 μW ~ 500 μW; the wavelength: 488 nm; CSU-LS2WF, Solution Systems, Japan) through a water immersion objective lens (ACROPLAN 63X, Zeiss, Germany). To align the larval ventral nerve cord (VNC) on the focal plane, the slide glass was put on an adjustable slope by moving a fulcrum. The recording duration was 233 seconds (700 frames). To suppress the noises, we applied smoothing processing with the method described in (Eilers, 2003), using a second-order penalty and *λ* = 1.5. We clipped a region of 100×100 pixels from one sample for the application of our patch-wise modularity-based clustering and non-negative matrix factorization (section 2.4) (Fig. 7.C).

In the following sections, imaging data of each focal plane is described by *l_txy_*(*l_txy_* = 0,1,…,2^14^ – 1), where *t* is the frame number and *x,y* are pixel coordinates (*t* = 0,1,…,*T* — 1, *x,y* = 0,1,…,*D* — 1, *T* = 700 frames, *D* = 512 px).

#### 2.2.2. Baseline correction and elimination of deformation and drift

The signal intensity at each pixel could be changed by two main factors: change in fluorescence intensity according to sensing calcium ions by the probe, and experimental artifacts, including bleach of the fluorophore of the sensors and drift of the sample. We needed to extract the former from the calcium imaging data, and the latter should be removed. Here we assumed that the artifacts changed slowly, and we could treat them as a change in the baseline of the signal. Therefore, the signal intensity at each pixel of the fluorescence movie was assumed to be the product of the baseline and the ratiometric change in probe signal intensity as follows:

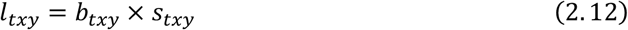

where *b_txy_* was the baseline, and *s_txy_* was signal-to-baseline ratio.

To suppress the effect of temporal change of the baseline, we extracted the component of *b_txy_* by applying baseline detection processing as described in (F. Zhang et al., 2020) with *λ* = 2 × 10^5^.

The extracted *b_txy_* was a fluorescence distribution without calcium transient (Fig. C.3.A). Therefore, *b_txy_* was first used to correct the sample deformation and drift. Each frame of *b_txy_* was transformed to match the specific frame (*b*_*t*_0_*xy*_, *t*_0_ = *T*/2; *T* = the total frame number) with a rigid transformation (homography), using cv2 function “findTransformECC” (Fig. C.3.B). Then, the baseline detection was applied again to the spatially transformed imaging data (*x,y* → *x*’,*y*’) and new baselines *b_tx’y’_* were calculated (Fig. C.3.C). Finally, the temporal profiles of calcium signals without baseline change were extracted (Fig. C.3.D). The pure neuronal signal *s_tx’y’_* was calculated by rearranging the Eq. (2.12):

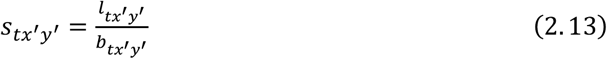

The primes of the spatial variables are omitted in the following sections: *s_txy_*.

#### 2.2.3. Extraction of signals from focal planes

The data *s_txy_* obtained above contained blurred signals from activities out of focus because the focal planes were placed in a three-dimensional object filled with calcium probes (Fig. 3.A). These blurred and coarse signals made it difficult to extract individual bouton activities. We formulated *s_txy_* as a summation of fine (well-focused) and coarse (out-offocus) components:

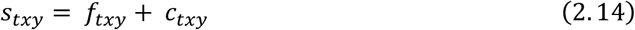

here *f* and *c* were fine and coarse components, respectively.

**Fig. 3.**
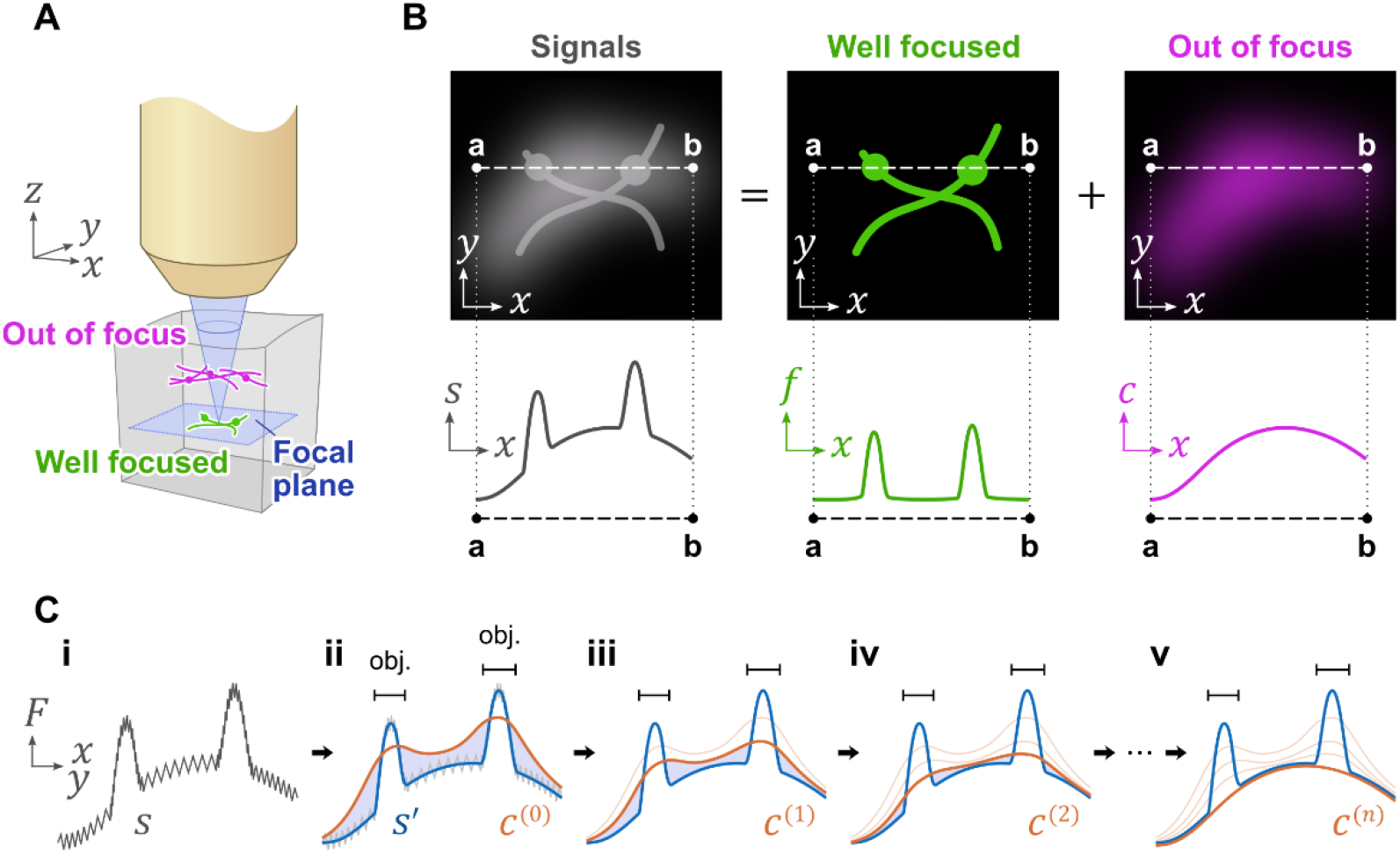
Baseline detection based on image contrast. (A) Schematics show that fluorescence signals contain fluorescence from both the focal plane and planes out of focus. (B) (Top): The detected signals (left) are the summation of signals from the focal plane (middle) and from out of the focal plane (right). (Bottom) Plot of signal intensities between points a and b in the corresponding top images. Note that the contrast of the signals in the well-focused image is high while those out of focus are low. (C) Principle of the two-dimensional baseline detection. The baseline is extracted by repeated calculation of a recurrence formula. The gray line in panel i is the original signal. Blue lines in panels ii to v indicate the smoothed signals, and orange lines show the candidates of its baselines. The signals from the objects on the focal plane (obj.) are extracted as the difference between the orange and blue lines by iteration. See section 2.2.3 for details.

The data *s_txy_* was decomposed into *f_txy_* and *c_txy_* based on the difference in the spatial scale of intensity change. In each data frame, the coarse component looked like a sntly changing baseline, and narrow peaks of the well-focused component were mounted ι that baseline (Fig. 3.B). Therefore, like the decomposition of *l_txy_* into *s_txy_* and *b_txy_* in action 2.2.2, the spatial structure of *f_txy_* and *c_txy_* could be segregated by utilizing the scale difference of intensity change (Fig. C.4.A). In this study, we applied two-dimensional bsseline detection to *s_txy_*, using Gaussian blur. Gaussian blur is defined as follows:

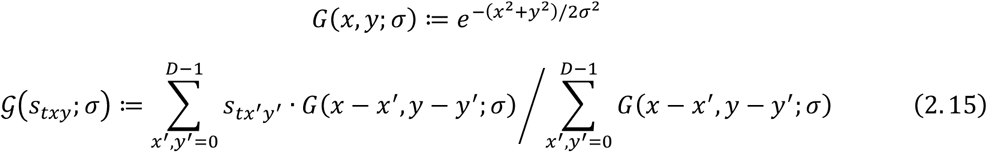

At first, Gaussian blur was applied to the extracted signals *s_txy_* (*σ*_1_ = 1) to reduce detailed fluctuation (Fig. 3.C.i):

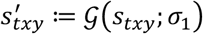

Then, the coarse component that corresponded to the baseline was calculated by the recurrence formula of 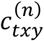:

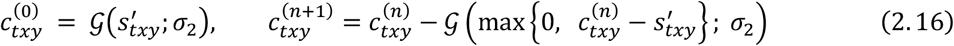

In this formula, we took advantage of the fact that when we set the standard deviation of the Gaussian filter between the scales of the coarse component and fine component, the coarse component was robust to the Gaussian filter while the fine component was sensitive. First, 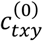 was obtained by applying the Gaussian filter to the original image 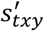 (Fig. 3.C.ii). Since the intensity of the signals was larger than that of the background, 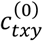 became smaller than 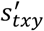 in the pixels that had significant signals from the well-focused plane (object region, hereafter. See horizontal bars in Fig. 3.C), but larger than 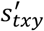 in the other region (non-object region, hereafter). Therefore, the sign of 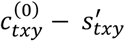 could be used to segregate the object region from the non-object region. By applying the Gaussian filter to the region where 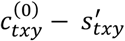 was positive, we could obtain the coarse component of the difference between 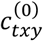 and 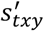 in the non-object region. By subtracting the coarse component of the difference from 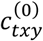 itself, a new coarse component 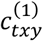 that had less difference from the original image 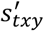 in the non-object region but didn’t have a fine component was obtained (Fig. 3.C.iii). By repeating these steps, 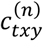 would converge to the curve that almost fit the original image 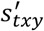 in the non-object region but didn’t have a fine component. The well-converged curve 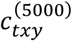 was treated as the baseline, and 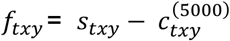 was used as signals from the focal plane.

The significance of this decomposition was observed in the comparison between activity waves spontaneously propagating in forward or backward directions on the VNC (Fig. C.4.B). It has been shown that several interneurons are activated in either forward or backward waves (Carreira-Rosario et al., 2018; Fushiki et al., 2016; Hiramoto et al., 2021; Kohsaka et al., 2019). In our data of pan-neuronal calcium recordings in the neuropil, such direction-specific activity could be observed only in the fine component (Fig. C.4.B). Therefore, the decomposition step was critical to extracting the proper activity of the neurons.

### 2.3. Artificial data of calcium signals

Artificial spatiotemporal data simulating calcium imaging were generated to test the performance of clustering methods. The dataset was made in three steps: building a 3-dimensional neuropil model with boutons, creating a calcium signal for each of the boutons in the 3-dimensional neuropil, and generating spatiotemporal signals in a 2-dimensional section from the 3-dimensional while adding realistic noises and perturbations.

#### 2.3.1. Building a 3-dimensional neuropil model with boutons

A group of boutons were built by splitting a 3-dimensional array of voxels (50 x 50 x 50 voxels) using the k-means algorithm with a fixed number of boutons. The x-y plane where *z* = 25 was treated as the focal plane in our artificial data (see section 2.3.3.) The total bouton number on the focal plane was set by tuning the total bouton numbers in the whole volume. The total bouton number in the volume was set to around 225, 450, 900 or 1,800 to let the bouton number on the focal plane be 25, 50, 100, or 200, respectively. First, each voxel was randomly assigned to one of the boutons. Then, the bouton assignment for all the voxels was updated five times by the k-means algorithm based on the Euclidean distance between voxels. In the end, each voxel was assigned to one bouton, and the whole 3-dimensional array was split into the pre-fixed number of boutons

#### 2.3.2. Creating calcium signals in the 3-dimensional neuropil model

Next, we generated a temporal signal to be assigned to each of the boutons. The length of the signals was 100, 200, 400, 800, and 1600 frames, and each of the temporal signals included multiple peaks. Each peak was modelled by a Gaussian function with time as a variable. By changing the temporal differences between the peaks of different bouton signals, two activity patterns were created: burst-like and sporadic activity. The former simulated a cooperative firing pattern seen in wave propagation, whereas the latter exhibited a less cooperative activity pattern. To simulate a peak, the time, width, and height of the peak were required. The time of the peaks in the signals was sampled from a Gaussian distribution, and the two different patterns (burst-like and sporadic activity) were made by setting a distinct deviation for the timing of signal peaks among the boutons. For the signal data of 100 frames, we set the mean of the peak times to be at the frames of 20, 55, and 80, and the standard deviation of them to be 2 frames for the burst-like activity and 10 for the sporadic activity. For the longer signal data of 200, 400, 800, and 1600 frames, the same samplings were repeated the required number of times (for example, 16 times for the data of 1600 frames) and concatenated in series. The width and height of the peaks were sampled from the Gaussian distribution (mean = 4 frames and standard deviation = 0.5 frames for the burst-like activity; mean = 2 frames and standard deviation = 0.5 frames for the sporadic activity) and the gamma distribution (shape index = 3 and scale index = 1/3), respectively. We set the baseline of the signal to be the same as the mean value of the gamma distribution 1.

#### 2.3.3. Simulation of realistic 2-dimensional imaging data

To simulate realistic calcium imaging data, we added several perturbations to the spatiotemporal signals generated above, mimicking the variation in the probe expression level, photobleaching, mixing of fluorescence signals from out of the focal plane, specimen drift and deformation, and random noises.

At first, to simulate the variation of the expression level of calcium probes in boutons, we sampled a relative probe level *b_n_* for the n-th bouton from the Gaussian distribution (mean = 2, standard deviation = 0.3). By multiplying the calcium signals generated above by the relative probe level *b_n_,* the signal considering the expression variation 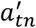 could be described as follows:

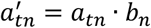

where *a_tn_* was the artificial signal of the n-th bouton at frame *t* (*t* = 0,1,…,*T* – 1; *T* = the total frame number).

Next, the photobleaching during the imaging was modelled by multiplying all the signals by the exponential decay factor:

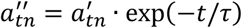

where *τ* = 1,000 frames was the decay time.

To mimic the mixing of fluorescence signals among different focal planes, blurred images out of the focal plane were superimposed on the image of the focal plane (*z*_focus_ = 25). Gaussian blur was applied to each image of the focal planes (*z* = 1,…,50) in the 3-dimensional array of voxels (50 x 50 x 50 voxels). The sigma for the Gaussian blur, which determined the size of the blurring filter, was obtained by *σ_z_* = (|*z* – *z*_focus_| + 1) · tab*α*, where *z* was the position of the plane along the z-axis (*z* = 0,.,49), *z*_focus_ = 25, sin*α* = NA/*n*_water_, NA = 0.9, and *n*_water_ = 1.333. Then, the blurred images, including that of the focal plane, were averaged by the weight of 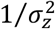:

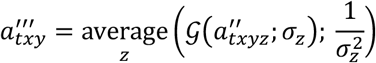

where 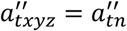 when (*x,y,z*) ∈ *n*-th bouton, and 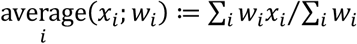.

The specimen drift and deformation were simulated by applying homography transformation to the image of the focal plane. The transformation could be uniquely determined by setting the destination of four corners of the image ((*x,y*) = (0,0), (*L*,0), (0,*L*), (*L,L*)), where *L* = 50. The transformation mapped the corners onto (*d*(*t*),*d*(*t*)), (*d*(*t*),*L* – *d*(*t*)), (*L* + *d*(*t*),—*d*(*t*)), (*L* + *d*(*t*),*L* + *d*(*t*)), where *d*(*t*) = 3(2*t*/*T* – 1) at frame *t*. The other points were mapped by interpolation. The signal intensity at each pixel in the transformed data was calculated by interpolation. This definition gave a drift in the x-direction, contraction of the side *x* = 0, and extension of the side *x* = *L*.

Finally, we added random noise sampled from a Gaussian distribution to each pixel at each frame of the data independently. The mean of the Gaussian distribution was zero, and the standard deviation *σ*_noise_ was chosen from 0.1-1.5 to change noise intensity. The noise level was comparable to that in biological calcium imaging data (Fig. C.5). The spatiotemporal data generated by these steps were used as the artificial imaging data. The x-y plane image where *z* = 25 in the data generated in section 2.3.1 was regarded as the correct configuration of boutons. The signal data assigned to each bouton built in section 2.3.2 was treated as the correct calcium signal. The performance of the clustering algorithms was assessed by comparing the outcome of the algorithms and the correct configuration or calcium signals.

### 2.4. Data analysis using CaImAn

Our algorithm was compared with CaImAn, an open-source tool for calcium imaging data analysis (version 1.9.9) (Giovannucci et al., 2019). Following the documentation of CaImAn, the analysis pipeline was built by modifying parameters in a released demo pipeline for one-photon imaging data decomposition (“demo_pipeline_cnmfE.ipynb”, See Table 1 for the parameter modification.) CaImAn was originally developed to extract cell bodies from calcium imaging data. To apply it to our bouton imaging data, the bouton size was used as the parameter of neuron size in CaImAn.

**Table 1.**
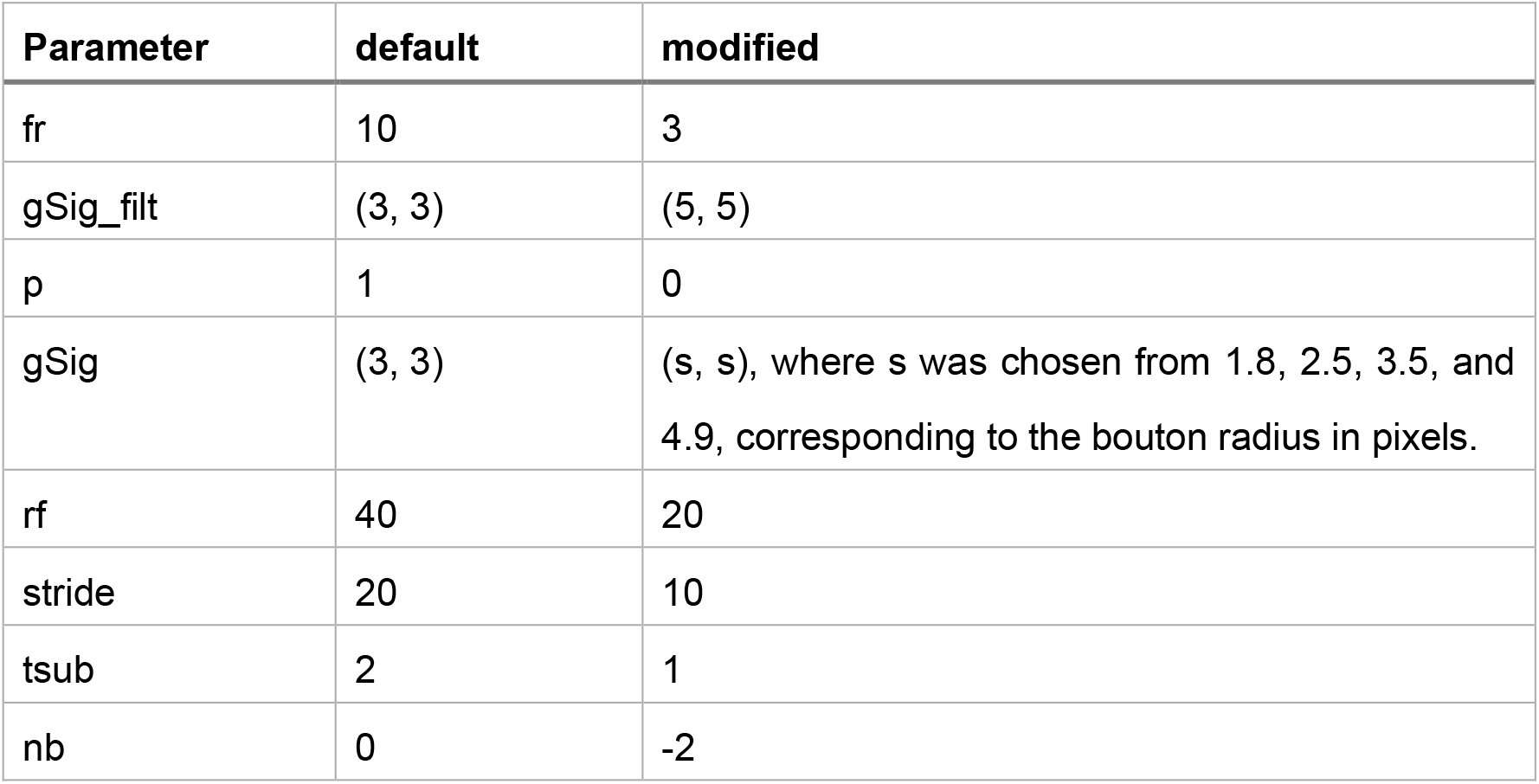
Parameter modification in the CaImAn demo pipeline.

| Parameter | default | modified |
| --- | --- | --- |
| fr | 10 | 3 |
| gSig_filt | (3, 3) | (5, 5) |
| p | 1 | 0 |
| gSig | (3, 3) | (s, s), where s was chosen from 1.8, 2.5, 3.5, and 4.9, corresponding to the bouton radius in pixels. |
| rf | 40 | 20 |
| stride | 20 | 10 |
| tsub | 2 | 1 |
| nb | 0 | -2 |

In the CaImAn pipeline, piece-wise rigid motion correction and spatiotemporal deconvolution based on a constrained non-negative matrix factorization (CNMF) (Pnevmatikakis et al., 2016) algorithm were performed. CNMF detected each bouton (spatial component) and extracted its activity (temporal component). The spatial components from CNMF described a bouton as a 32-bit float image. To compare the output from CNMF with the one from our algorithm, the 32-bit images were binarized by Otsu’s thresholding method (Otsu, 1979). The CaImAn pipeline had an option to use a deconvolution algorithm to estimate spike trains under the assumption that spike trains were sparse. However, since the activity patterns in the fly larval motor circuits were not sparse but burst-like, the deconvolution algorithm was not applied.

### 2.5. Evaluation of clustering performance

The performance of clustering methods was assessed by evaluating spatial and temporal accuracy in signal detection. The spatial accuracy was measured by using the Dice coefficient. Let 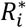 be the set of pixels included in the *i*-th true bouton, and *R_j_* be the set of pixels included in the *j*-th detected bouton-like region. The Dice coefficient between 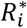 and *R_j_* is defined as follows:

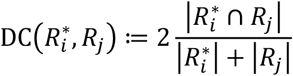

Where |·| indicates the number of pixels included in the set. 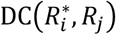 indicates similarity of 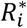 and *R_j_*, taking a value between 0 and 1. To measure the spatial accuracy of the bouton detection, we defined spatial similarity (*A_s_*) by a weighted average of maximum Dice coefficient with the weight of the pixel number in boutons:

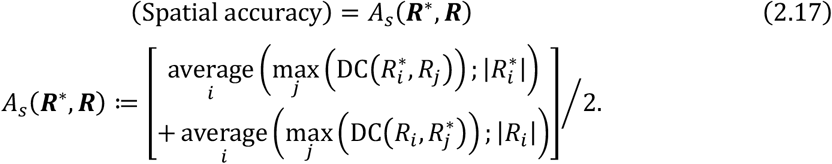

Note that *A_s_*(***R***^(**1**)^***R***(**2**)) = *A_s_*(***R***^(**2**)^,***R***^(**1**)^) and 0 ≤ *A_s_* ≤ 1. *A_s_*(***R***^(**1**)^***R***^(**2**)^) = 1 if and only if ***R***^(**1**)^ and ***R***^(**2**)^ were identical. The weight of the average (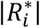 or |*R_i_*|) were imposed because the Dice coefficient was not stable when either region was small.

To measure the temporal accuracy of activity extraction, we defined temporal similarity (*A_t_*) between the true signal of the *i*-th true bouton 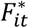 and the extracted signal of the *j*-th detected bouton *F_jt_*, which had the maximum Dice coefficient with the *i*-th true bouton. The Pearson correlation coefficient between 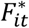 and *F_jt_* in terms of *t* was calculated. The average of the correlation coefficient over all the true boutons was used as the temporal accuracy:

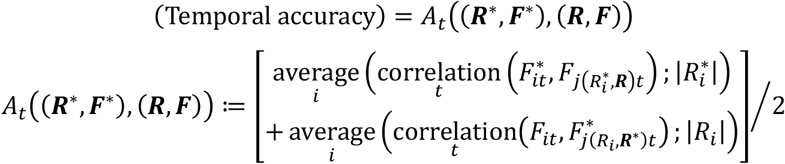

where 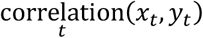 is the Pearson correlation coefficient between *x_t_* and *y_t_* in terms of *t*, 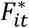 was the true signal of the *i*-th true bouton, *F_jt_* was extracted signal of the *j*-th detected bouton and 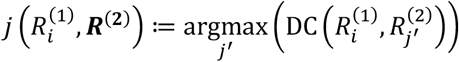. Note that 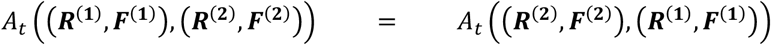 and −1 ≤ *A_t_* ≤ 1. *A_t_*((***R***^(**1**)^,***F***^(**1**)^),(***R***^(**2**)^,***F***^(**2**)^) = 1 if and only if (***R***^(**1**)^,***F***^(**1**)^) and (***R***^(**2**)^,***F***^(**2**)^) were identical except for scaling and offset of ***F***.

Our algorithm used random processes to optimize the configuration of pixel clusters. Therefore, the results could differ between data processing trials with the same data. To test the consistency of the signal extractions among trials, we calculated a consistency measure defined as follows:

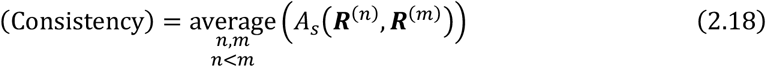

where 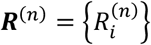 was the set of detected bouton-like regions from the *n*-th trial. The consistency from ten trials for each artificial data of distinct parameters was calculated (Fig. 5).

### 2.6. Evaluation of calculation time

To measure calculation duration, we applied the bouton detection to artificial data with frame numbers 100, 200, 300 and 400, while the pixel number was fixed to 2,500 (50 x 50). (Fig. 9.A-D). In addition, the calculations were applied to artificial data with pixel numbers 1,024 (32 x 32), 2,500 (50 x 50), 3,969 (63 x 63), 5,476 (74 x 74) and 7,056 (84 x 84), while the numbers of the true boutons were proportional to the pixel numbers (41, 100, 159, 219 and 282), and the frame number was fixed to 100 (Fig. 9.E-H). The artificial data was generated as the burst-like signals with *σ*_noise_ = 0.1.

## 3. Results

### 3.1. Performance assessment of the Patch-wise modularity-based clustering with artificial data

We established a novel method, Patch-wise modularity-based clustering (“PQ-based clustering” hereafter), to cluster bouton-like structures from calcium imaging data (See section 2.1 for details.). In short, to cluster pixels in a calcium imaging movie into a set of boutons, a similarity between a pair of pixels was defined based on their activity patterns. Under the assumption that each bouton was spatially compact, the similarity was calculated between pixels within a pre-determined distance to obtain a patch-wise similarity matrix. Then, a set of boutons in the image were extracted by optimizing the clustering of pixels to let the pixels within each cluster have high similarity. To assess the performance of the PQ-based clustering, spatial accuracy (the bouton configuration) and temporal accuracy (the calcium signal) were analyzed (See section 2.5.)

To this aim, we generated artificial calcium imaging data. The artificial data were designed referring to the actual calcium imaging data from the neuropil of the larval motor circuits. The neuropil consisted of densely packed boutons and exhibited propagation activity, a progression of local burst-like activity along the body axis. To model these features, an array of voxels (50 x 50 x 50) was divided into hundreds of boutons (Fig. 4.A), and cooperative burst-like activity was assigned to the boutons (Fig. 4.B) (See sections 2.3.1 and 2.3.2 for details.) In addition, several perturbations were added to the artificial data to mimic real calcium imaging data, considering the variation in the probe expression level, photobleaching, mixing of fluorescence signals from out of the focal planes, specimen drift and deformation, and random noises (Fig. 4.A’ and B’) (See section 2.3.3 for details.)

**Fig. 4.**
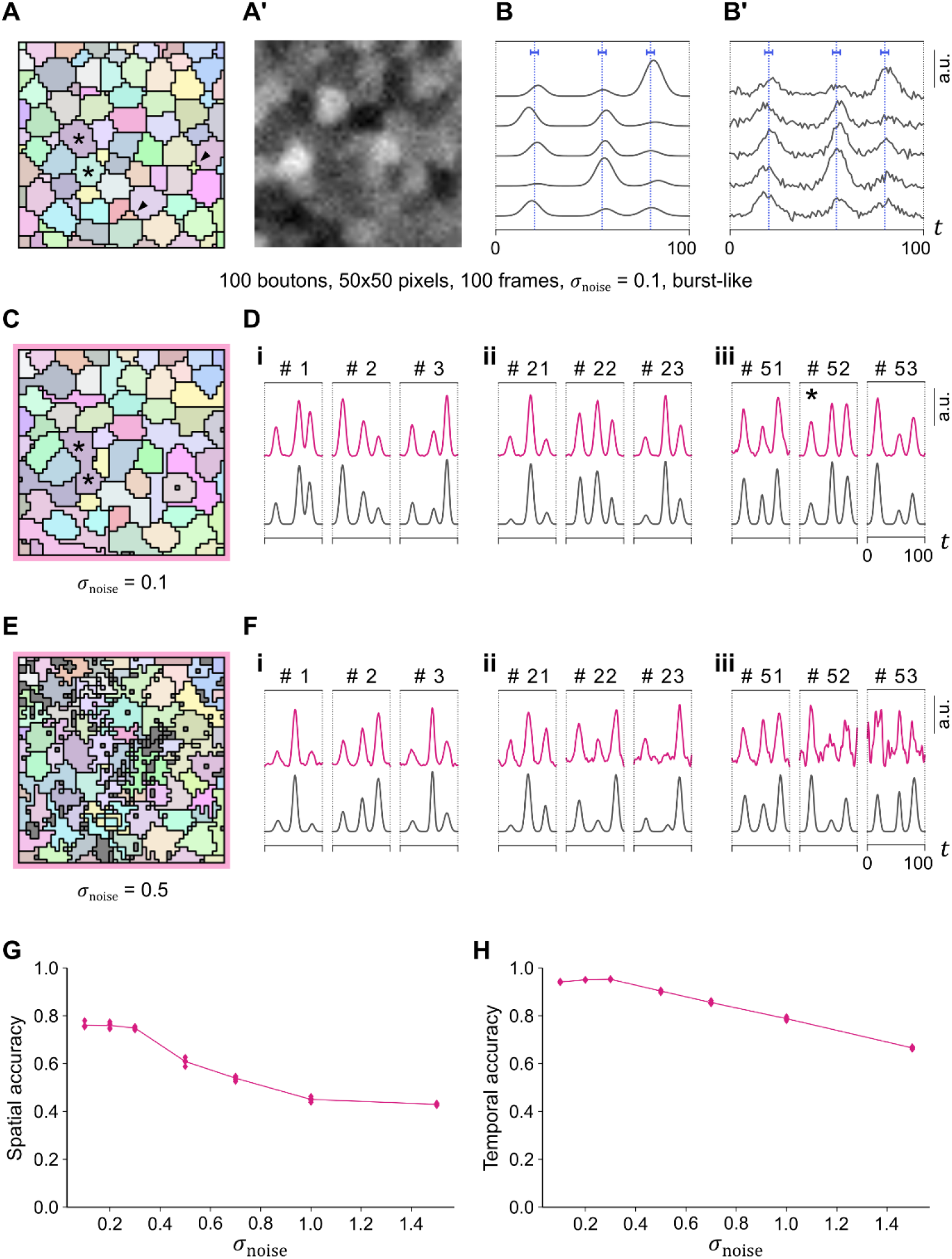
PQ-based clustering successfully detected boutons and calcium signals from burstlike artificial data. (A) Correct configuration of boutons in the artificial data. One hundred boutons were included in the 50 x 50 pixel array. Different colors indicate each bouton. Arrowheads indicate small boutons that the clustering could not recognize. (A’) Signal intensity in one frame in the burst-like artificial data. (B) Correct signals of five example pixels. (B’) Several perturbations were added to the correct signals to achieve realistic artificial data. The traces show the signal in five pixels selected in (B) with the perturbations under the condition of *σ*_noise_ = 0.1. Vertical blue lines in (B) and (B’) indicate the center of each burstlike event, and horizontal blue bars in (B) and (B’) indicate the standard deviation of peak timings around each center. (C-F) Bouton extraction from the burst-like artificial data with *σ*_noise_ = 0.1 (C and D) and *σ*_noise_ = 0.5 (E and F). (C and E) Detected bouton configuration by PQ-based clustering. The color of each cluster corresponds to that of its closest correct bouton in (A). Stars in (A) and (C) indicate boutons that the clustering could not discriminate. (D and F) Extracted signals (magenta traces) by PQ-based clustering from the data shown in (C) and (E), respectively, and the correct signals (gray traces). Pairs of correct and detected boutons with large Dice coefficient (i: top 1-3, ii: top 21-23, iii: top 51-53) are shown. The pair of the 52^nd^ Dice coefficient (star) is the merged boutons indicated by stars in (C). (G and H) Spatial (G) and temporal (H) accuracy of PQ-based clustering of the burst-like artificial data of each noise intensity.

**Fig. 5.**
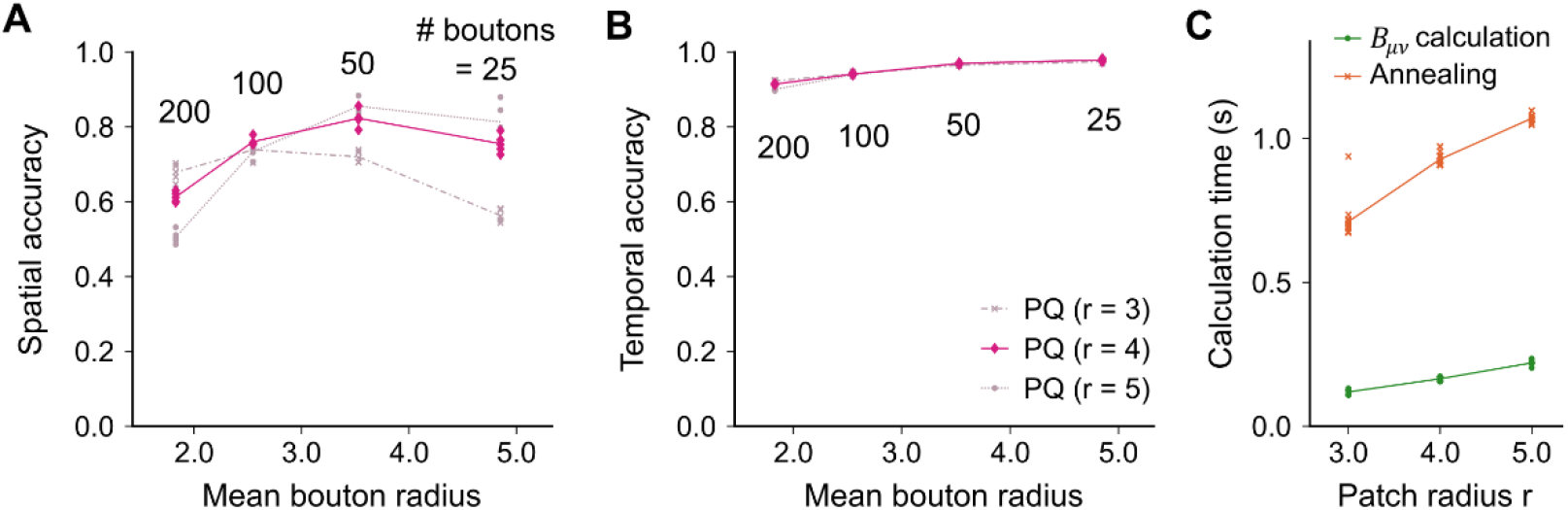
Detection accuracy could be optimized by tuning the patch radius r. (A and B) Spatial (A) and temporal (B) accuracy of PQ-based clustering with a different patch radius, r = 3 (gray crosses), 4 (magenta diamonds), and 5 (gray circles). (C) Calculation time for PQ-based clustering with a different patch radius r. Green dots and orange crosses indicate the calculation time for *B_μν_* and simulated annealing, respectively.

#### 3.1.1. Performance of the PQ-based clustering method

We applied PQ-based clustering to the artificial data to test the signal extraction performance. From the data of the burst-like activity with a moderate noise (*σ*_noise_ = 0.1), the PQ-based clustering method extracted most of the boutons (Fig. 4.C). The signal from the extracted boutons was consistent with its correct calcium signal (Fig. 4.D). It should be noted that some small boutons that were less than 10% of the average size could not be detected (arrowheads in Fig. 4.A). And when two adjacent boutons had similar temporal activities, they were extracted as a single bouton (compare stars in Fig. 4.A and C). Although some pixels were mislabeled (< 30%), the PQ-based clustering extracted the majority of the boutons from the artificial dataset. Even when the noise level in the artificial data was increased up to *σ*_noise_ = 0.5, the majority of the boutons and consistent calcium signals were still extracted (Fig. 4.E and F). If the noise level increased to *σ*_noise_ = 1, which corresponded to the condition when the noise intensity was comparable to that of calcium signals, PQ-based clustering could detect only a half of the boutons (Fig. 4.G) but still with high temporal accuracy (Fig. 4.H). Accordingly, PQ-based clustering could extract the calcium signals from boutons in the artificial dataset mimicking real calcium imaging data. In particular, the spatial accuracy was high when the noise level was lower than the signal intensity.

#### 3.1.2. The adjustability of extracted bouton size in the PQ-based clustering method

In the PQ-based clustering, the patch radius r, which determined the range of distance between pixels to calculate the similarity, was a key hyper-parameter and could influence the clustering performance. This effect was tested by analyzing the results of the PQ-based clustering with a distinct patch radius (r = 3, 4, and 5 pixels). When the data whose mean bouton radius was 1.8 pixels, the PQ-based clustering with a patch radius of 3 gave the highest spatial accuracy. On the other hand, for the data with an average bouton radius of 4.9, the PQ-based analysis with r = 5 provided the best spatial accuracy (Fig. 5.A). Temporal accuracy was not significantly affected by the difference in the patch radius (Fig. 5.B). Because patch radius r determined the range of pixel pairs to be calculated, the calculation time of the clustering was increased by the upregulation of r. However, the increased calculation time was maintained at a manageable level (Fig. 5.C). Therefore, the patch radius could adjust the spatial scale in the PQ-based clustering. In the following analyses, the artificial data with an average bouton size of 2.5 pixels and the patch radius r = 4 were used unless otherwise noted

#### 3.1.3. Consistency of the PQ-based clustering method

The optimization step in the PQ-based clustering contained random processes (see section 2.1.3.) Therefore, distinct trials could give different results even if the data was identical. To check the consistency of the outcome of the PQ-based clustering, we compared the results among multiple trials with the same artificial imaging data (For the measure of consistency, see section 2.5.) The consistency was about 0.8 or more for every case and kept at about 0.9 if the noise level was low (Fig. 6.A and B). Accordingly, even though random processes were included, the PQ-based clustering method could provide consistent results over multiple trials.

**Fig. 6.**
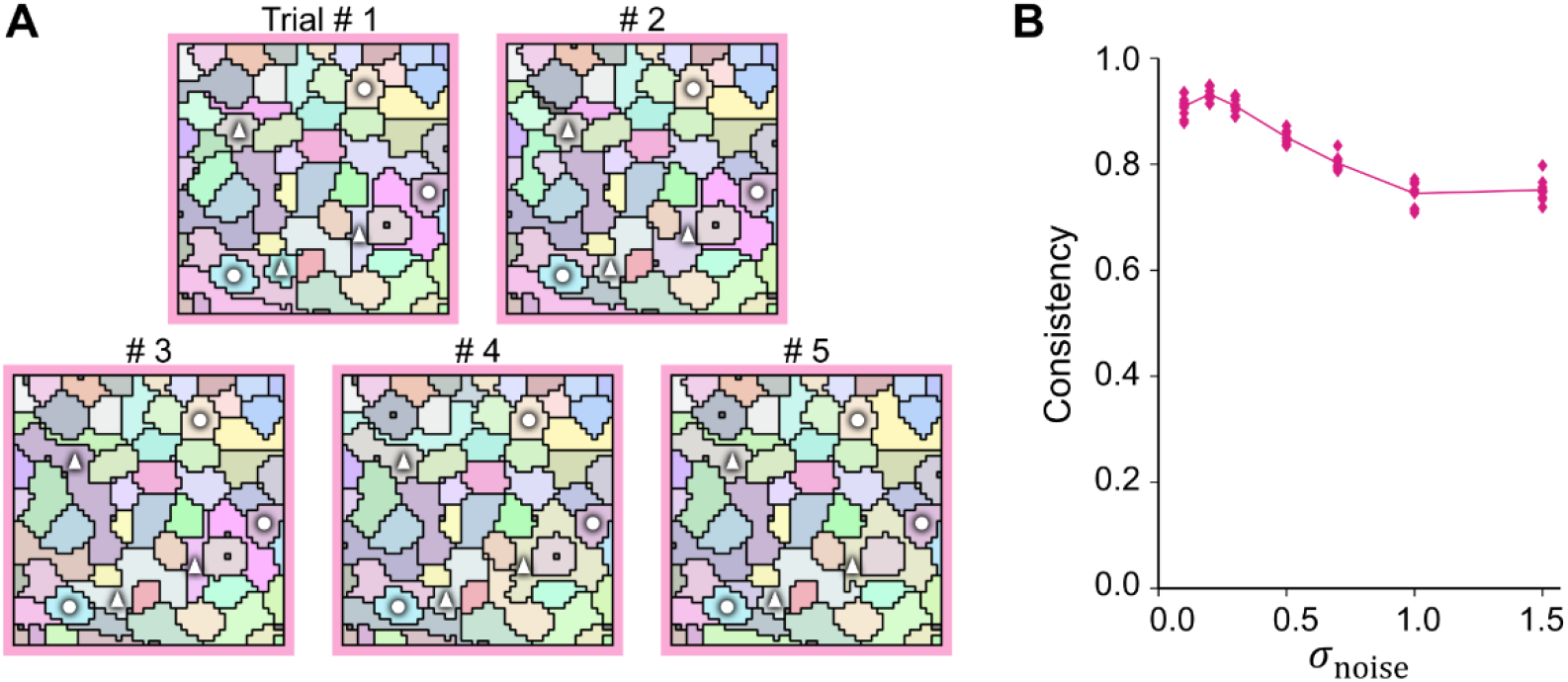
PQ-based clustering provided consistent results over multiple trials. (A) Detected bouton configuration of five trials from the burst-like artificial data with *σ*_noise_ = 0.1. The color of each cluster corresponds to that of its closest correct bouton shown in Fig. 4.A. White circles and triangles mark the same clusters among the five trials. The circles indicate regions with relatively high spatial consistency, while the triangles indicate regions with relatively low spatial consistency. (B) Spatial consistency was quantified as the average spatial similarity among five trials (see section 2.5 for details.) Each data point indicates the spatial similarity of a pair of trials.

### 3.2. Performance comparison with the CaImAn algorithm

We next compared the PQ-based algorithm with CaImAn (Giovannucci et al., 2019), one of the standard tools to extract cell bodies from calcium imaging data (see section 2.4 for details.)

#### 3.2.1. Detection accuracy in the analysis of the burst-like activity data

The CaImAn algorithm detected cell bodies under the assumption that neurons exhibited sparse firing activity but not cooperative burst-like activity. We tested the CaImAn algorithm’s performance to analyse our artificial data with burst-like activity. From the data with moderate noises (*σ*_noise_ = 0.1), CaImAn could detect most correct boutons (Fig. 7.A). However, the spatial accuracy was lower than that of the PQ-based clustering (Fig. 7.E). This was because CaImAn extracted some clusters that didn’t correspond to any boutons whose number was about the total number of the correct boutons. One reason for this mislabeling would be that the nature of cooperative activity made CaImAn difficult to find the boundary of some boutons. In the temporal aspect, some peaks were not detected in CaImAn (Fig. 7.B), and the temporal accuracy of CaImAn was lower than that of the PQ-based clustering (Fig. 7.F). This might be caused by CaImAn’s ability to decompose a mixture of activity signals from spatially overlapped components: the correct boutons were not overlapped in our artificial data, but CaImAn possibly assumed that some activity peaks came from other overlapped boutons and eliminated them from the activity extraction. Therefore, some peaks would be removed. When the noise level was increased up to *σ*_noise_ = 0.5 and more, the overall bouton configuration and spatial accuracy were maintained (Fig. 7.C and E). However, the noises strongly affected the extracted signals, and temporal accuracy decreased (Fig. 7.D and F). Overall, PQ-based clustering could detect bouton configurations and calcium signals more accurately for the artificial burst-like activity data than the CaImAn algorithm.

**Fig. 7.**
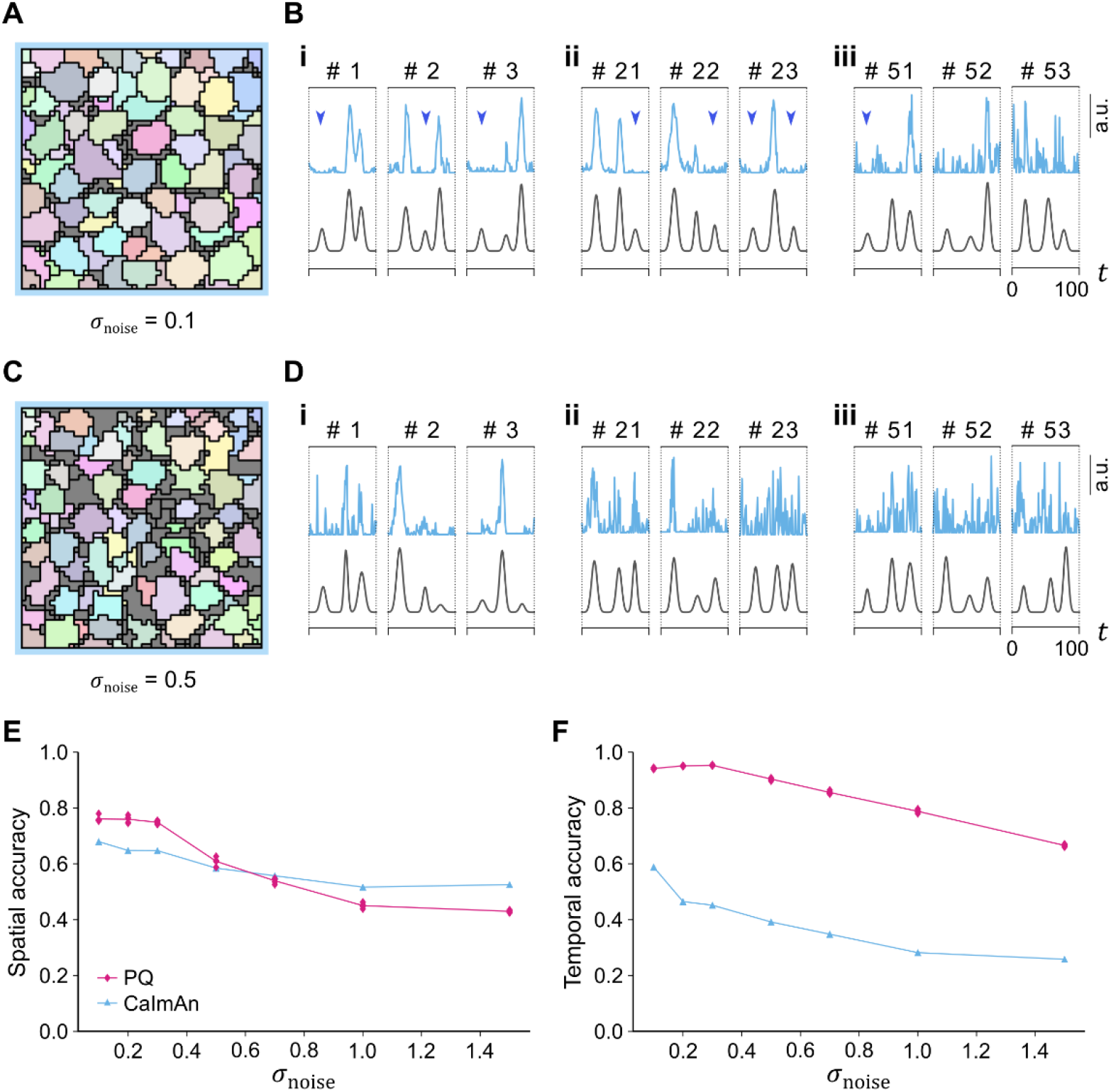
Comparison of PQ-based clustering and CaImAn in the analysis of the burst-like artificial data. (A-D) Bouton extraction from the burst-like artificial data with *σ*_noise_ = 0.1 (A and B) and *σ*_noise_ = 0.5 (C and D). (A and C) Detected bouton configuration by CaImAn. The color of each cluster corresponds to that of its closest correct bouton shown in Fig. 4.A. (B and D) Extracted signals (cyan traces) by CaImAn from the data shown in (A) and (C), respectively, and the correct signals (gray traces). Pairs of true and detected boutons with large Dice coefficient (i: top 1-3, ii: top 21-23, iii: top 51-53) are shown. Blue arrowheads indicate the timings of peaks missed in the results by CaImAn. (E and F) Spatial (E) and temporal (F) accuracy in the analysis of the burst-like artificial data of each noise intensity by PQ-based clustering (magenta diamonds) and CaImAn (cyan triangles). The plot of PQ-based clustering in (E) and (F) are the same as in Fig. 4.G and H, respectively.

#### 3.2.2. Detection accuracy in the analysis of sporadic activity data

The observation above implied the low detection accuracy attributed to the nature of burst-like activity patterns. To pursue this possibility, we generated artificial data with sporadic activities and applied the two clustering algorithms to them (see section 2.3 for generating the sporadic activity data.) CaImAn extracted peaks in correct signals more robustly, drastically improving its temporal accuracy (compare Fig. 8.F and H with Fig. 7.B and F.) These results suggested that the PQ-based clustering should be useful to extract bouton configurations and calcium signals, especially from calcium imaging data with burst-like activities.

**Fig. 8.**
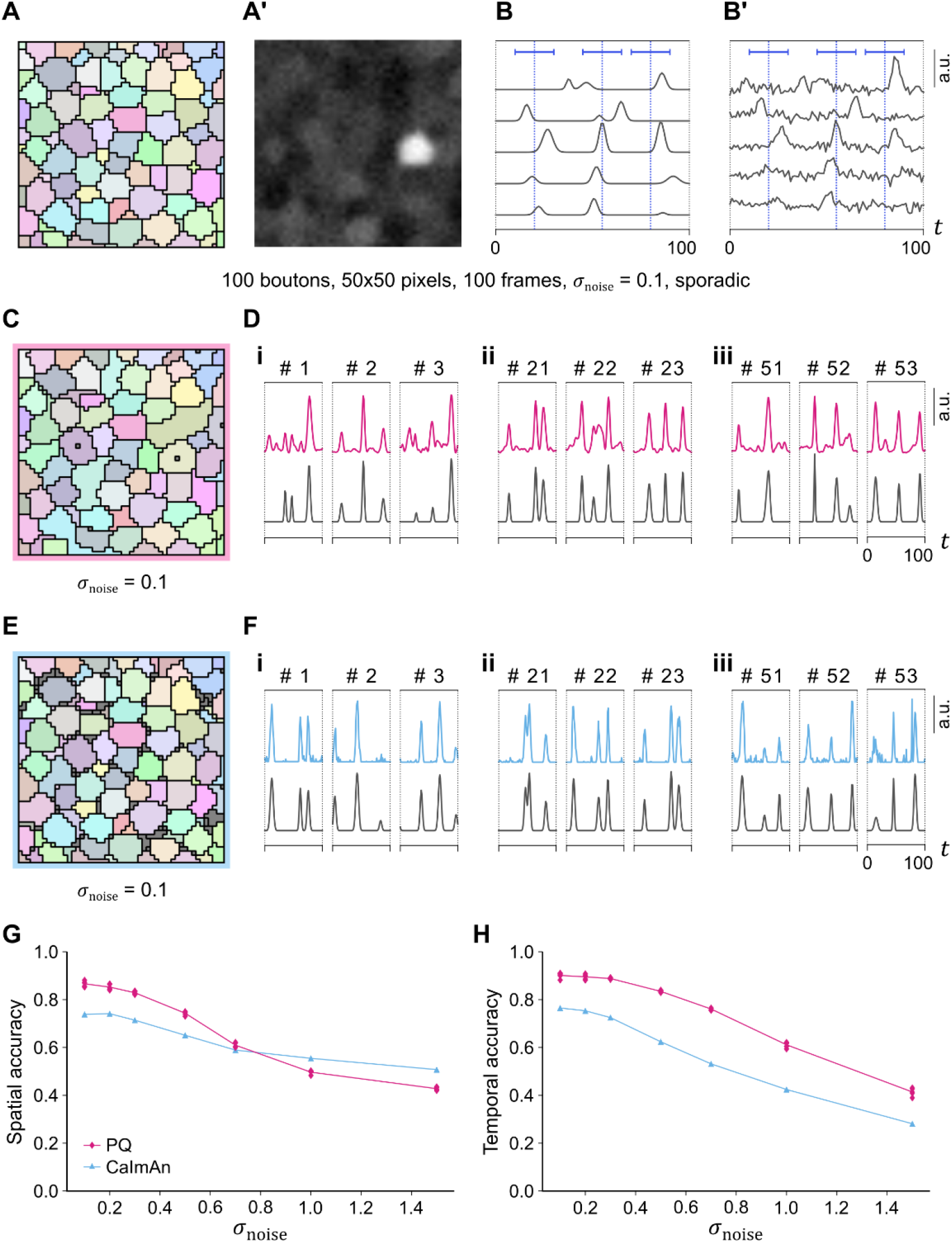
Both PQ-based clustering and CaImAn successfully detected boutons and calcium signals from sporadic artificial data. (A) Correct configuration of boutons in the artificial data. This data is the same as in Fig. 4.A. (A’) Signal intensity in one frame in the sporadic artificial data. (B) Correct signals of five example pixels. (B’) Several perturbations were added to the correct signals to achieve realistic artificial data. The traces show the signal in the five pixels selected in (B) with the perturbations under the condition of *σ*_noise_ = 0.1. Vertical blue lines indicate the center of each sporadic event, and horizontal blue bars indicate the standard deviation of peak timings around each center. (C-F) Bouton extraction from the sporadic artificial data with *σ*_noise_ = 0.1 by PQ-based clustering (C and D) and CaImAn (E and F). (C and E) Detected bouton configuration of PQ-based clustering and CaImAn, respectively. The color of each cluster corresponds to that of its closest correct bouton in (A). (D and F) Extracted signals by PQ-based clustering (magenta traces in D) and CaImAn (cyan traces in F) from the data shown in (C) and (E), respectively, and the correct signals (gray traces). Pairs of correct and detected boutons with large Dice coefficient (i: top 1-3, ii: top 21-23, iii: top 51-53) are shown. (G and H) Spatial (G) and temporal (H) accuracy in the analysis of the sporadic artificial data of each noise intensity by PQ-based clustering (magenta diamonds) and CaImAn (cyan triangles).

**Fig. 9.**
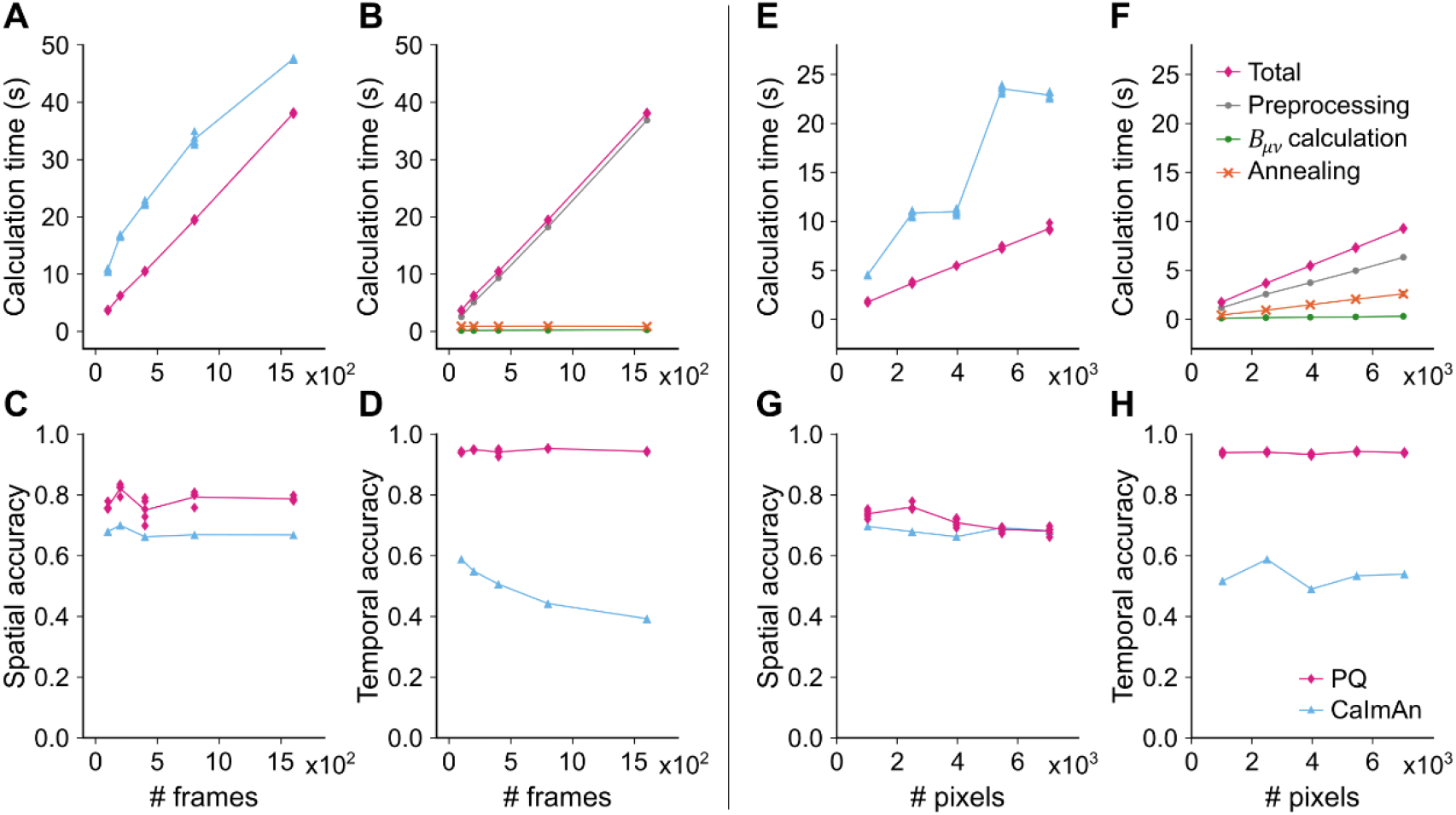
Comparison of the calculation time between the PQ-based clustering and CaImAn. (A-D) Performance of the PQ-based clustering and CaImAn in the analysis of burst-like artificial data of various total frame numbers. (A) Calculation time. (B) The breakdown of (A). (C) Spatial accuracy. (D) Temporal accuracy. (E-H) Performance of the PQ-based clustering and CaImAn in the analysis of burst-like artificial data of various total pixel numbers. (E) Calculation time. (F) The breakdown of (E). (G) Spatial accuracy. (H) Temporal accuracy. In (A), (C), (D), (E), (G), and (H), magenta diamonds indicate the results of PQ-based clustering, whereas cyan triangles indicate those of CaImAn. In (B) and (F), magenta diamonds, gray circles, green circles, and orange crosses represent the total calculation time for PQ-based clustering, the time for preprocessing, the time to calculate the modularity matrix *B_μν_,* and the time for simulated annealing.

#### 3.2.3. Calculation time

The calculation time was one of the critical factors in the data analysis. The time required to conduct clustering on a standard desktop computer with the PQ-based clustering and CaImAn was analyzed (see section 2.6 for details.) In processing the 100-frame data, the calculation time for the PQ-based clustering was shorter than that for the CaImAn algorithm (Fig. 9.A). To test this observation in different data sizes, we generated artificial data with a different frame or an image size. First, the frame number was changed from 100 to 1600. While the calculation time for both algorithms increased as the number of frames increased, the time for PQ-based clustering was shorter in the range tested while keeping better detection accuracy of the PQ-based clustering (Fig. 9.A, C, and D). Next, the image size was changed from 1000 to 8000 pixels. Calculation time for both CaImAn and PQ-based clustering increased linearly as the number of pixels went up, and the calculation duration for PQ-based clustering was about a half or one-third of that for CaImAn while keeping better detection accuracy of the PQ-based clustering (Fig. 9.E, G, and H). Consequently, the PQ-based clustering required less calculation time and provided more accurate clustering in the range of data size we tested.

The time for PQ-based clustering consisted of three parts: preprocessing, calculating the patch-wise modularity matrix, and simulated annealing. Of these three, preprocessing occupied the longest time in the processing time. On the other hand, calculation of the patch-wise modularity matrix and simulated annealing accounted for less time (Fig. 9.B and F). Accordingly, efficient computing in the patch-wise calculation that drastically reduced the number of the calculation of the similarity between pixels would underlie the fast bouton detection by the PQ-based clustering.

### 3.3. Performance assessment with experimental data

To assess the performance of the clustering method with experimental data, we performed pan-neuronal calcium imaging of the neuropil of the fly VNC and applied preprocessing, such as baseline correction, the elimination of deformation and specimen drift, and the extraction of signals from the focal plane, to the data (see Methods section 2.2.) We obtained spatiotemporal data (512 x 512 pixels, 700 frames) from three isolated CNS (Fig. 10.A and A’). In each imaging data, the propagations of neural activity waves were observed multiple times.

**Fig. 10.**
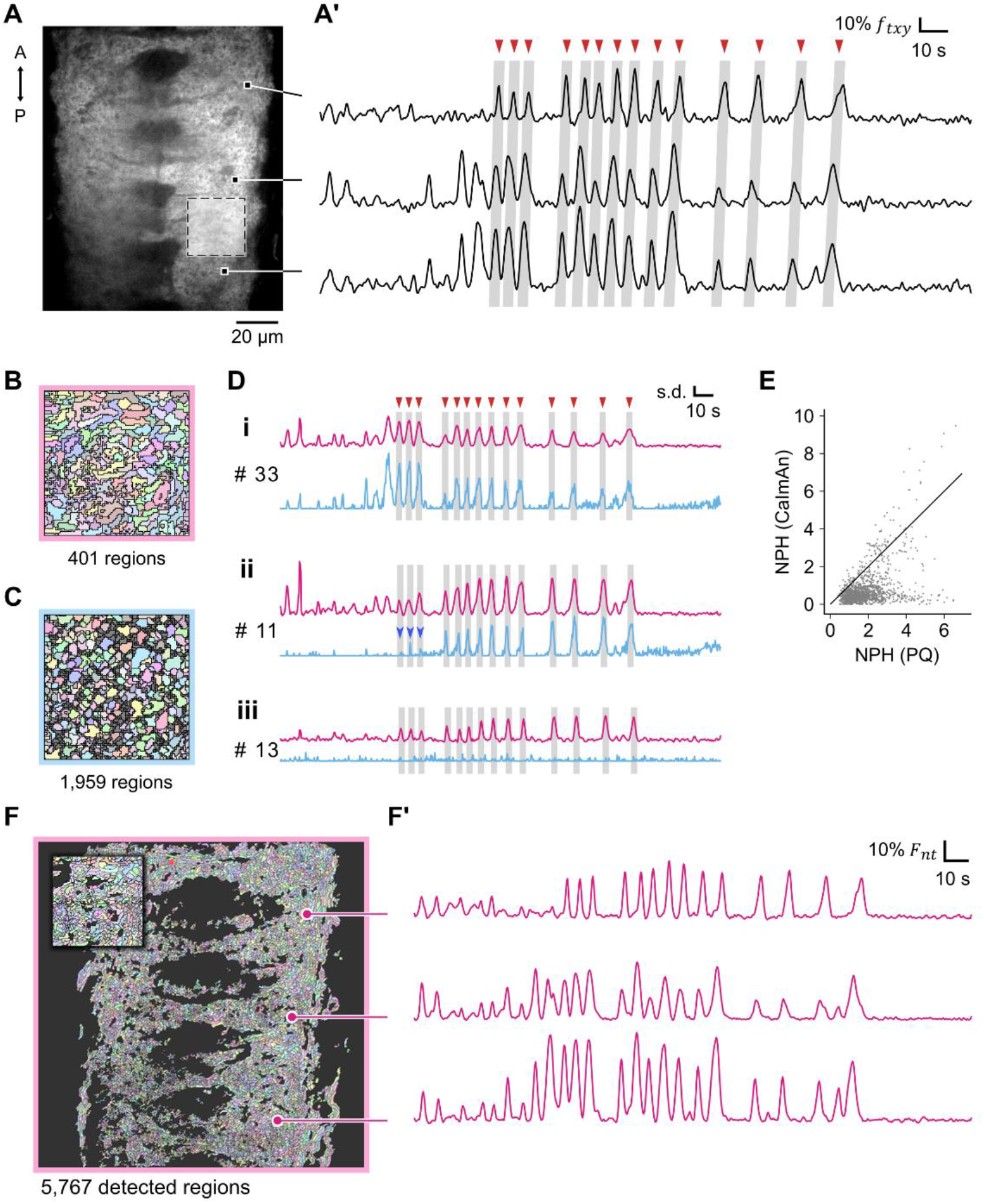
PQ-based clustering was applicable to real calcium imaging data. (A) Snapshot image of calcium imaging of the isolated CNS of a *Drosophila* larva. Dashed square indicates the position where the image was clipped for (B) and (C). (A’) Preprocessed temporal traces of example pixels in (A). Red arrowheads and shaded areas indicate activity propagations from the posterior to anterior segments corresponding to fictive crawling behavior. (B and C) Clustering results of the clipped data by the PQ-based clustering (B) and CaImAn (C). Each detected cluster in (B) is indicated by a different color. For each cluster in (B), a cluster in (C) that overlapped with the cluster in (B) the most was filled with the same color. Clusters in (C) that were not painted based on the overlap with clusters in (B) were filled with gray. (D) Extracted temporal signals from the clipped data using PQ-based clustering (magenta) or CaImAn (cyan). Three pairs of detected clusters from the two methods with a large Dice coefficient (i: top 33, ii: top 11, iii: top 13) are shown. Blue arrowheads indicate the timings of peaks missed in the results by CaImAn. Red arrowheads and shaded areas indicate the same activity propagation events in (A’). The signal intensity obtained by each method was normalized by the standard deviation (s.d.) of peak heights of all the detected clusters in the indicated activity propagation events (PQ-based clustering or CaImAn). (E) Comparison of the peak heights of spatially matched 110 clusters from PQ-based clustering and CaImAn. Each data point represents normalized peak heights (NPHs) of a matched cluster. The vertical axis shows NPH by the PQ-based clustering, while the horizontal axis shows that by CaImAn. The black line indicates NPH (PQ) = NPH (CaImAn), where the normalized peak height by the PQ-based clustering is the same as that by CaImAn. (F) Detected bouton-like regions in the whole imaging data of the same sample by PQ-based clustering. Each of the bouton-like regions is shown by a different color. The upper-left window shows an expanded image of the region indicated by the white dotted box in (A). (F’) Temporal traces of the example bouton-like regions in (F) taken from the same positions in (A). Note that this clustering was done without manual tracing and showed consistent results with (A’).

First, we compared the performance between PQ-based clustering and CaImAn by image data of small size. A region of 100×100 pixels from one sample was clipped (Fig. 10.A, the square of dashed lines). Both the PQ-based clustering and CaImAn could detect bouton-like regions from the calcium imaging data. The size of bouton-like structures extracted by the PQ-based clustering was consistent with that by CaImAn (the PQ-based clustering: 0.6 ± 0.4 μm (n = 401), CaImAn: 0.7 ± 0.2 μm (n = 1,959)). However, since CaImAn allows the overlapping of boutons, the total numbers of the extracted boutons were larger in the CaImAn output than in the PQ-based clustering result (Fig. 10.B and C). Furthermore, while the total areas of the extracted boutons in the PQ-based clustering results were the same as the total area of the original image, those in the CaImAn results were 4 times larger than the original image (400 ± 10%), which implied that the results by CaImAn should include a large number of clusters that didn’t correspond to correct boutons. To investigate the signals from the bouton-like structures extracted by the two clustering methods, we matched the clusters from the two algorithms based on the overlapped areas measured by the Dice coefficient and compared the signals of matched clusters (Fig. 10.D). Some matched boutons exhibited consistent activity between the two (Fig. 10.D.i). However, as observed in the analysis of the artificial data (Fig. 4.D and 7.B), some peaks were detected by the PQ-based clustering but not by CaImAn (Fig. 10.D.ii and iii). Therefore, the peak heights of the activity of the bouton-like structures were higher in the result of the PQ-based clustering than that of CaImAn (Fig. 10.E). These observations suggest that the PQ-based clustering can detect peaks of bouton activity more reliably than CaImAn.

Next, we applied PQ-based clustering to the full-size imaging data and extracted bouton-like regions (4,950 ± 640 (mean ± std., n = 3)) and their temporal profiles (Fig. 10.F and F’). The calculation for the clustering of the full-size data using PQ-based clustering took 40.6 ± 0.6 (mean ± std., n = 3) minutes (preprocessing: 93%, calculation of *B_μν_*: 2%, simulated annealing: 4%). Accordingly, PQ-based clustering allowed us to extract bouton-like structures from experimental calcium imaging datasets.

### 3.4. Network analysis among the extracted bouton-like regions

Finally, as an application of the clustering data, we attempted to extract a network structure of the fly CNS based on the clustering results. We used the correlation of calcium signals between two bouton-like structures to define the interaction between them. We calculated the correlation coefficient between every bouton pair’s temporal profiles, binarized them with a threshold of 0.9, and obtained an unweighted graph of bouton-like structures and their connections (Fig. 11.A).

**Fig. 11.**
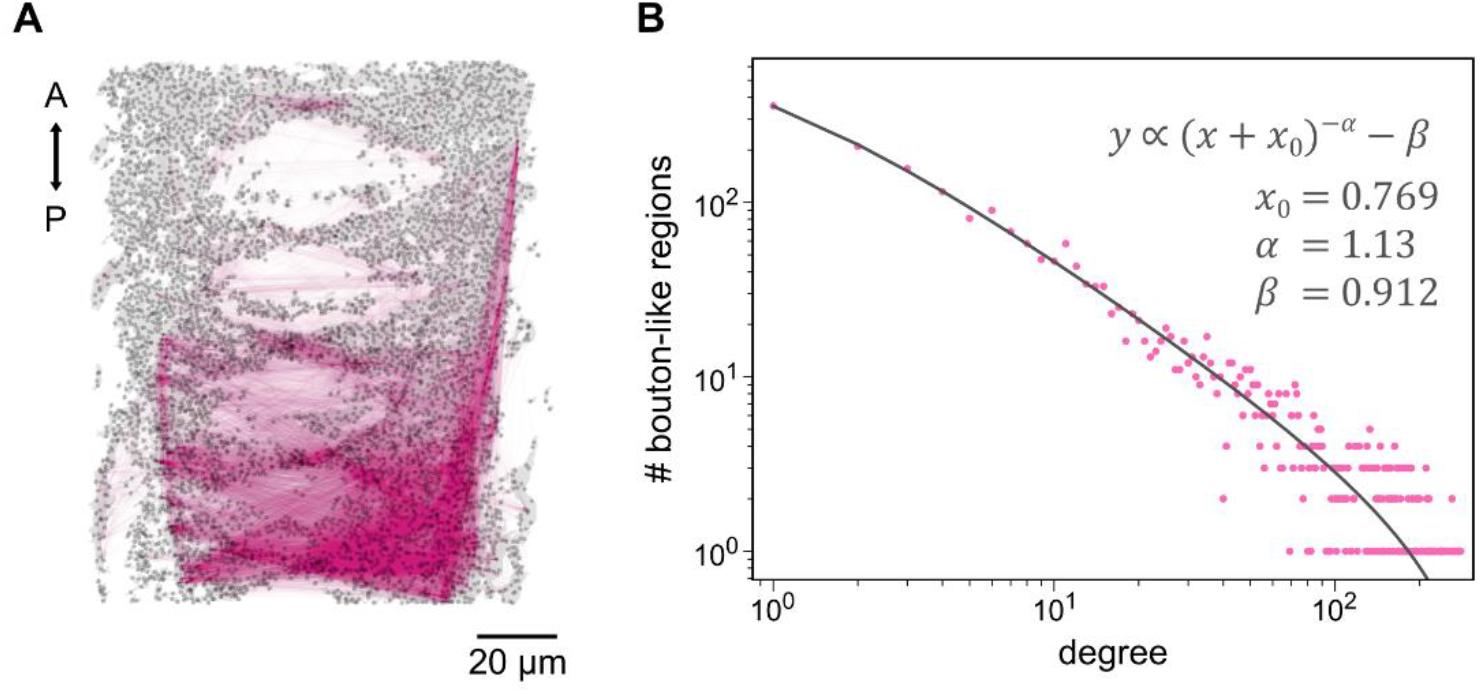
Correlation networks among activities of bouton-like regions formed scale-free networks. (A) Extracted network structure of the same sample of the CNS of a *Drosophila* larva in Fig. 10. Each black dot indicates the centroid of each detected region. Each magenta line connects two regions with a correlation coefficient larger than 0.9. (B) Histogram of degrees in the network of neural activity in the CNS of *Drosophila* larvae. Magenta dots show the network’s degree distribution in (A) in a log-log plot. The horizontal axis indicates the degree of each bouton-like structure, while the vertical axis indicates the number of boutonlike regions. The black line fits the histogram whose model is shown in the panel.

Network analysis of the graphs suggested they were scale-free (Barabási & Albert, 1999). We found that the distribution of degrees, which were the numbers of edges from bouton-like structures, followed the power law (Fig. 11.B). The scaling exponent *α* was 1.14 ± 0.01 (mean ± std., n=3), which implied that there existed hub nodes in the networks (Cohen et al., 2000; Pastor-Satorras & Vespignani, 2001; Wang et al., 2011). This was consistent with previous studies that had shown functional connectivity in the human brain forms scale-free graphs (Eguíluz et al., 2005; Stam & Reijneveld, 2007; van den Heuvel et al., 2008).

## 4. Discussion

Here, we introduced a new methodology to detect bouton-like regions from artificial and experimental calcium imaging data. To detect bouton-like regions from images where boutons were densely concentrated, we made progress in three aspects of algorithms and theories: first, we established pre-processing methods to extract signals from focal planes in experimental imaging data by using a two-dimensional baseline detection algorithm; second, we defined patch-wise modularity, which is an expanded version of Newman and Girvan’s modularity (Newman & Girvan, 2004), and mathematically proved its validity and availability; third, we engineered an algorithm to maximize the patch-wise modularity based on simulated annealing method (Kirkpatrick et al., 1983).

The first progress was to extract signals from well-focused planes in the imaging data. Since the neurites in the neuropil are densely packed in a three-dimensional space, imaging data for population activities in the neuropil typically include fluorescence signals from out of the focal plane. In this study, we extracted signals from the well-focused plane by a two-dimensional baseline detection algorithm. Several algorithms have been proposed to determine baselines of one-dimensional spectral data, such as Raman and infrared spectra, based on a wavelet transform (Bertinetto & Vuorinen, 2014; Qian et al., 2017; Shao & Griffiths, 2007), derivative methods (Leger & Ryder, 2006; Schulze et al., 2005), polynomial fitting (Gan et al., 2006; Lan et al., 2007; Lieber & Mahadevan-Jansen, 2003; Zhao et al., 2007), and penalized least squares (Baek et al., 2015; He et al., 2014; F. Zhang et al., 2020; Z. M. Zhang et al., 2010). However, these methods were not readily expanded to the two dimensions because their implementation in higher dimensions could be complicated. In our approach, we introduced iterative Gaussian blur to realize baseline determination in two-dimensional images and successfully detected baseline from experimental calcium imaging data.

The second achievement was the introduction of patch-wise modularity. Although modularity is a widely used quality measure to detect community structures in a network (Bruno et al., 2015; Chen et al., 2009; Fine et al., 2006; Humphries, 2011; Sporns & Betzel, 2016; Torre et al., 2019), it has been criticized because the size of its detected communities is affected by the size of the whole network (Fortunato & Barthélemy, 2007; Lancichinetti & Fortunato, 2011; Rosvall & Bergstrom, 2013). This “resolution limit problem” arises because the definitions of the modularity involve the term the total strength of all edges. In our calcium imaging data analysis, the resolution limit was about at least nine pixels, which hindered the detection of bouton-like structures whose size was about five pixels. Moreover, our targeting networks of similarity among pixels were large and dense (basically, all node pairs were connected), so it was almost impossible to calculate the total strength. Several researchers have reported that the resolution limit problem can be eliminated by multi-resolution modularity (Arenas et al., 2008; Reichardt & Bornholdt, 2006) or an edge weighting scheme (Cao et al., 2016; Haq et al., 2019; Lu et al., 2018). However, these methods could not be applied to our analysis because they still needed to calculate the total strength. Or other methods tackled the resolution limit problem by using quality measures without such global summation of edge strength (Li et al., 2008; Muff et al., 2005). However, these measures were also inapplicable to our case because they were efficiently calculated only when edges were sparse enough. Our patch-wise modularity was defined without global quantities. Moreover, the pixel pairs contributing to the modularity estimation were restricted within each pixel-centered circle patch. Therefore, the calculation cost was within the acceptable level even with our large networks. In addition, since arbitral diameters of the patches were allowed in this measure, the virtual network size around each pixel could be easily modified. The flexibility in size enabled us to adjust the preferred size of pixel clusters obtained by optimizing the patch-wise modularity. Thanks to this measure, we succeeded in extracting bouton-sized pixel clusters independent of the resolution limit. The adjustability of the typical size of clusters (Fig. 5) might support the accessibility of structures in different scales from that of boutons in the neuropil, including cell bodies.

There is one limitation in using our method when decomposing a network. In our application of the detection of bouton-like structures, we took advantage of the fact that there is a stereotyped scale in boutons in the CNS. Using this bouton scale, we set the weight matrix as a binary function described in Eq. (2.5). In general applications, network decomposition methods are used to simultaneously detect communities of various sizes (Lancichinetti & Fortunato, 2011; Li et al., 2008; Rosvall & Bergstrom, 2013). The patch-wise modularity used in this paper cannot be applied to detect clusters with different sizes in the current form. However, the patch-wise modularity can be extended to meet this demand by setting other forms of the weight factor instead of Eq. (2.5). This factor is adjustable according to the purpose of each network decomposition, and we gave a mathematical proof of the validity of using any form of the weight matrix in the patch-wise modularity framework (Appendix B). Therefore, some modified version of the patch-wise modularity might detect various sizes of communities simultaneously in real-world networks by setting biases properly.

Finally, we developed a maximization algorithm for the patch-wise modularity based on the simulated annealing method. While there were several existing algorithms to maximize modularity (Clauset et al., 2004; Duch & Arenas, 2005; Guimerà et al., 2004; Newman, 2004, 2006; Reichardt & Bornholdt, 2006; Torre et al., 2019; White & Smyth, 2005), we made much account of the speed in the calculation, since the similarity networks among the imaging pixels were large and complex: about 2 x 10^5^ nodes and 7 x 10^6^ edges even after restricted within the patches, and 10^4^ clusters to be detected. Most maximization methods require calculation costs proportional to at least quadratics of these numbers, making it challenging to manipulate our data. The simulated annealing method can search parameter space at a fast speed (Metropolis et al., 1953). Furthermore, this method was suitable for our application: In the simulated annealing methods, the assignment of clusters was iteratively changed during the maximization process. Since boutons had continuous shapes, each pixel should belong to boutons in its neighbors. Moreover, we permitted duplicated identities for different and spatially segregated clusters since we could easily split them afterwards while the number of clusters to calculate could be reduced. These all properties contributed to the drastic reduction in the range of the search space and calculation costs. Thanks to these optimizations, the algorithm achieved the calculation costs proportional to simply the pixel number, with the fixed radius of the patches and number of iterations.

This study showed that our patch-wise modularity-based pixel clustering algorithm provided better results in the detection of bouton-like structure from neuropil data with burstlike activity than CaImAn (Giovannucci et al., 2019), which was based on constrained non-negative matrix factorization (CNMF) (Pnevmatikakis et al., 2016). Two properties of the neuropil data could be considered as its causes: 1. Temporal overlap of peaks between different boutons, and 2. Dense configuration of boutons.

In the burst-like activity data, signals of boutons that were close to each other exhibited cooperative activation (Fig. 4.B and 10.A’). This property was challenging for the NMF-based algorithm because NMF attempted to extract spatial components showing mutually different signals, which didn’t allow nearby boutons to exhibit similar activity. This issue could be found in the case of other methods to decompose calcium imaging data. For example, several decomposition methods based on principal component analysis, independent component analysis, similarity analysis, and singular value decomposition (Mölter et al., 2018; Mukamel et al., 2009; Shibue & Komaki, 2020) assume that the activities of different neurons are highly independent. In addition, CNMF deconvolves calcium signals into spike trains by promoting the sparseness of spikes. However, the neural spikes in our calcium imaging data were not sparse but showed coordinated burst activity related to motor control.

The dense configuration of boutons in the neuropil imaging data could also affect the clustering performance of CaImAn. CNMF is supposed to be used to extract cellular components that are dispersed in the neural tissue. In addition, various imaging data decomposition algorithms are based on the assumption that there is a background space between cell bodies. For example, an NMF-based algorithm by (Maruyama et al., 2014) includes background components, and the activity-based level set segmentation (ABLE) (Reynolds et al., 2017) detects regions on the premise that the target regions are surrounded by background regions. In contrast, boutons were densely packed in the neuropil images, and there was no space between boutons, which would make CNMF and other decomposition algorithms hard to segregate individual boutons. Furthermore, while clusters extracted by the PQ-based method were not spatially overlapped, CaImAn and other NMF-based methods allowed clusters to be overlapped. This property could underlie the observation that excess boutons were detected by CaImAn (Fig. 10.C).

It is important to note a limitation of our clustering method. Pixel clusters obtained by our methods can include not only a whole single bouton but also a fragment of a single bouton or a group of boutons. Because our method fixes the typical size of a cluster, a bouton larger than that scale might be split into multiple parts, or smaller boutons might be grouped into a single bouton. Actually, in the simulation with the artificial data, our method could not detect small boutons compared with patch radius r = 4.0 pixels and merged them with adjacent boutons (compare Fig. 4.A and C.) Our algorithm might group spatially overlapping boutons if the focal plane has a non-negligible thickness because distinct boutons with different depths can merge onto the same pixel. While some previous algorithms, including NMF, can decompose a pixel into multiple overlapping components (Maruyama et al., 2014; Mukamel et al., 2009; Pnevmatikakis et al., 2016; Shibue & Komaki, 2020), our pixel clustering method distributes exactly one cluster identity to each pixel. Our simulation showed the case where spatially overlapping boutons were grouped into a single cluster (Fig, 8.D): a part of detected boutons in high spatial accuracy exhibited additional activity peaks that were not included in the correct signals but were originated from spatially overlapped neighboring boutons.

Nevertheless, our method provides vast accessibility to the spatiotemporal population activity in the neuropil. Based on the extracted activity data, we could analyze network properties in the motor circuits (Fig. 11). Therefore, the spatiotemporal population activity of boutons could be a novel approach to examining neural network dynamics at the cellular level.

## Acknowledgements

We thank Bloomington Drosophila Stock Center for providing fly lines. We thank Dr. Ken Nakae for the critical comments on this paper. We also thank Drs. Dohjin Miyamoto, Suguru Takagi, Atsuki Hiramoto, Yingtao Liu, and Xiangsunze Zeng for informative discussions. This work was supported by MEXT/JSPS KAKENHI grants [17K19439, 19H04742, and 20H05048 to A.N. and 17K07042 and 20K06908 to H.K.].

## Appendix A: Patch-wise modularity

In one of the standard procedures for the analysis of modularity, for a given network whose connectivity strength between nodes *μ* and *v* is given by *A_μν_* (*A_μν_* = *A_νμ_* ≥ 0), the modularity matrix is defined as follows:

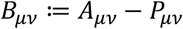

where *P_μν_* = *d_μ_d_v_*/*m*, *d_μ_* = ∑_*v*_*A_μν_*, *m*:=∑_*μ*_*d_μ_* = ∑_*μ,ν*_*A_μν_*. When we assume *p_μ_* = *d_μ_*/*m* is the probability distribution of connections linked to *μ*, it is clear that *P_μν_* = *m*·*p_μ_p_ν_* is the expected connectivity strength of arbitral networks with the same degree distribution *d_μ_* (note that *∑_ν_P_μν_* = *∑_ν_A_μν_* = *d_μ_*.)

In this study, we impose a constraint on the link between two nodes based on their spatial proximity: two nodes separated by more than five pixels are not linked. This condition cannot be met by the conventional assumption *P_μν_* = *d_μ_d_ν_*/*m* since it doesn’t use the information about their position. Accordingly, instead of defining *P_μν_* as being proportional to the product of *d_μ_*/*m* and *d_v_*/*m*, we assume that *P_μν_* should have non-zero components only if d(*μ,ν*) < *r* (See Eq. (2.5) in the Method section) and follow the same distribution of degrees as that of *Â_μν_*:= *W_μν_* · *A_μν_*. By this assumption, we can write the null network as follows:

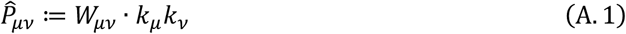

Here, *W_μν_* is defined in Eq. (2.5) and *k_μ_* is a positive vector that is chosen to satisfy 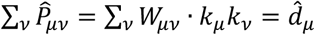. For each given *W_μν_* (not only the one defined in Eq. (2.5)) and 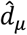, there always exists one and only one *k_μ_* (Theorem 1, Appendix B). 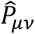 can be computationally given by using Theorem 2 in Appendix B. We define a patch-wise modularity matrix 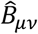 by this null network.

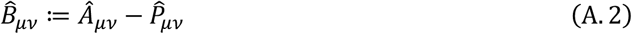

Because 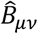 reduces to the original modularity matrix in case of *W_μν_* = 1 for every *μ,ν* (then 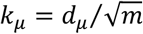), the definition gives an expansion of the conventional modularity. For a given cluster configuration *S* = {*s_μ_*}, the modularity is defined as follows:

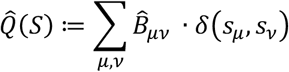

## Appendix B: Theorems and proofs

### Definition 1

Let *μ* and *v* be integers in {1,2,…,*N*} where 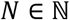.

*W_μν_* is a *N × N* non-negative symmetric matrix with positive diagonal components:

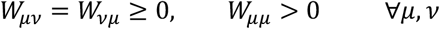

*k_μ_* and *d_μ_* are *N*-dimensional positive vectors:

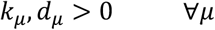

### Definition 2

For an *N*-dimensional vector 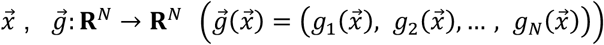 is defined as follows:

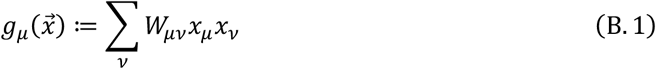

where 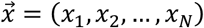.

### Theorem 1

For given *W_μν_* and *d_μ_*, there always exists one and only one *k_μ_* satisfying the following equations:

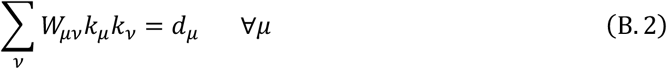

**Proof:**

### Lemma 1

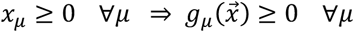

**Proof:**

By the assumption and Definition 1, all of the terms in 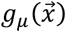 are positive. So, this statement is true.

### Lemma 2

For a given *μ*_0_ ∈ {1,2, …,*N*}, 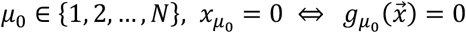

**Proof:**

Sufficiency: 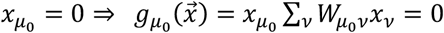

Necessity: 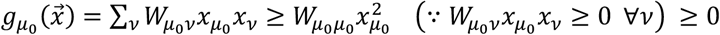

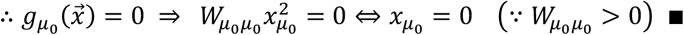

From Lemma 1 and 2, 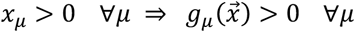. Therefore, when 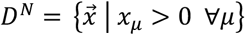, the mapping 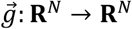 includes a partial mapping 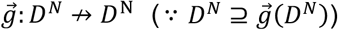. Because Theorem 1 is equivalent to the existence of the inverse mapping 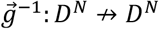, we attempt to prove 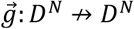 is bijective.

### Lemma 3

The partial mapping 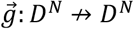 is injective.

**Proof:**

Lemma 3 is proved by contradiction.

Assumption 1: 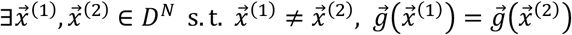

By using a line 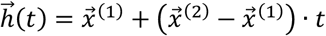, which passes through the two points 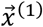 and 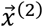, a scalar function 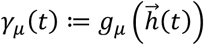 is defined. From Definition 2, *γ_μ_*(*t*) is described as a polynomial of *t*, which is at most quadratic. Therefore, from 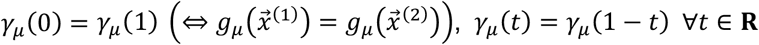 has to be satisfied for every *μ*. Accordingly, the following equation is required:

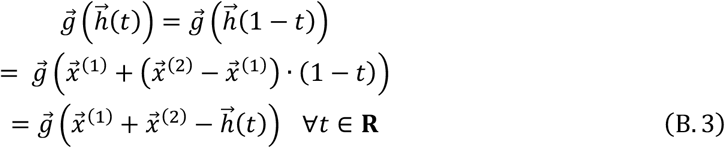

Here, there is at least one *μ*_0_ which satisfies 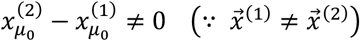. When 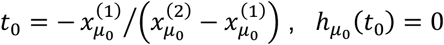 is valid. Then, the following equations are derived:

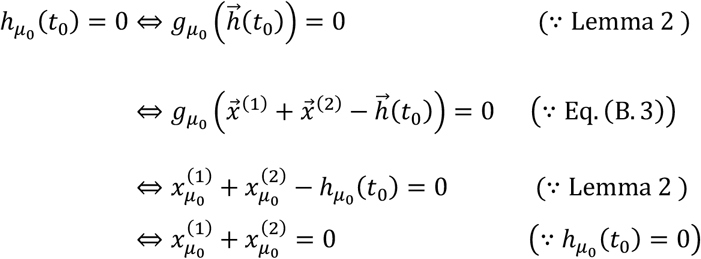

However, this is inconsistent with 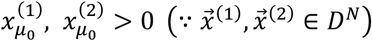. Therefore, Assumption 1 cannot be realized, and lemma 3 is true.

### Lemma 4

The partial mapping 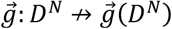 is a homeomorphism.

**Proof:**

Because the range is limited to the image of 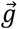, the partial mapping 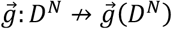 is surjective. Also, the mapping is injective from Lemma 3. Therefore, 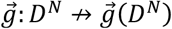 is bijective. From Definition 2, both 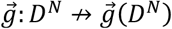 and 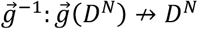 are continuous. Accordingly, the statement is true.

### Lemma 5

The partial mapping 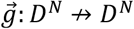 is surjective.

**Proof:**

Lemma 5 is proved based on mathematical induction.

Case 1: *N* = 1

For all *y* > 0, there exists *x* > 0, which satisfies 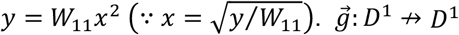 is surjective.

Case 2: *N* > 1

Assumption 2: 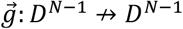 is surjective.

The boundary of *D^N^* is described as the following:

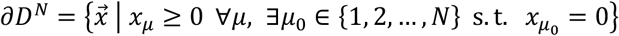

From Assumption 2 and Lemma 2, 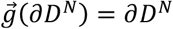. From Lemma 4, 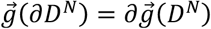. Therefore, 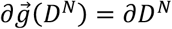. Then, from Lemma 4 and 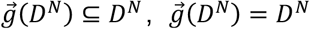 is derived. This indicates 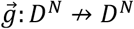 is surjective.

From Lemma 3 and 5, the partial mapping 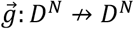 is shown to be bijective. This proves the validity of Theorem 1.

### Definition 3

A matrix sequence 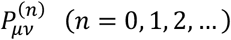 is defined by the following recurrence formula:

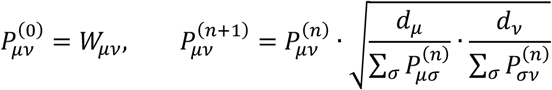

For 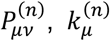 is defined as follows:

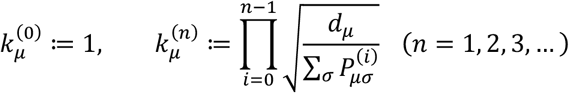

### Theorem 2

For given *W_μν_* and *d_μ_*, the positive vector *k_μ_* which satisfies Eq. (B. 2) is composed by the following:

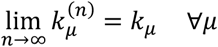

**Proof:**

### Lemma 6

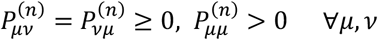

**Proof:**

Lemma 6 is proved based on mathematical induction.

Case 1: *n* = 0

From 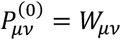 and Definition 1, 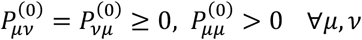

Case 2: *n* > 0

Assumption 3: 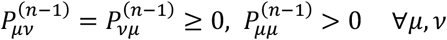

Obviously, 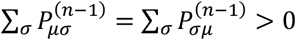. By combining with *d_μ_* > 0,

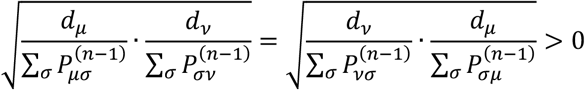

From 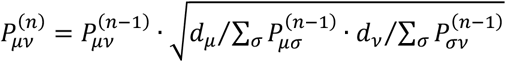 and Assumption 3,

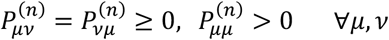

From Cases 1 and 2, Lemma 6 is valid for all *n* ∈ {0,1,2,…}.

From Definition 3 and Lemma 6,

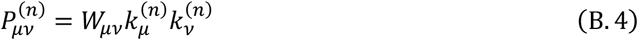

### Lemma 7

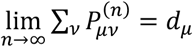 for all *μ* is a sufficient condition to satisfy Theorem 2.

**Proof:**

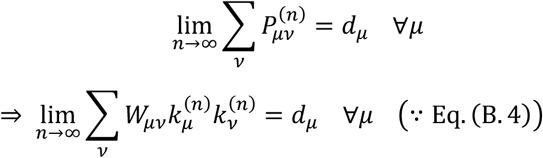

From Lemmas 4 and 5, the partial mapping 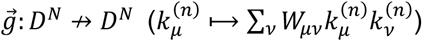 is a homeomorphism. Because *k_u_* is defined as the vector satisfying Eq. (B. 2), 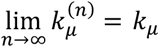 for all *μ*.

Because of Lemma 7, by using 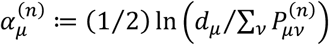, we attempt to show the following equation (*ln* is the natural logarithm):

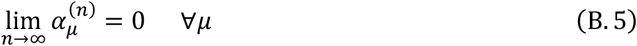

From Definition 3, 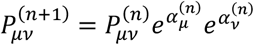 (*e* is Napier’s constant).

By substituting the right side for 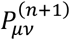 in 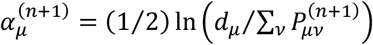, the following equation is obtained:

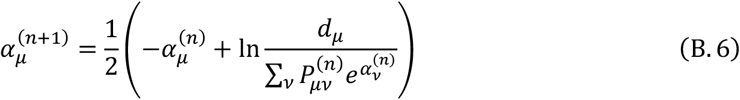

Here, because of 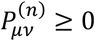, the following inequality is satisfied 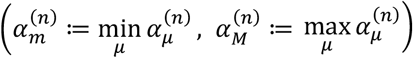:

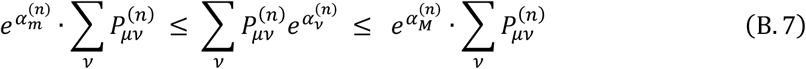

Therefore, from Eq. (B. 6), the following inequalities are derived:

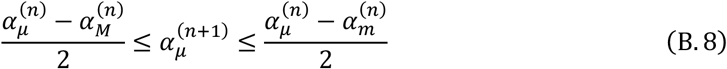

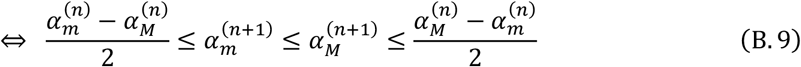

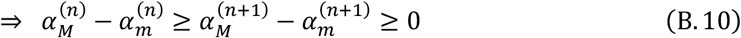

From Inequalities (B. 10), there is a non-negative finite value *ω* which satisfies 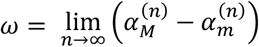.

Because 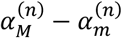 monotonously decreases according to *n*, 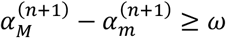.

Therefore, from Inequalities (B.9),

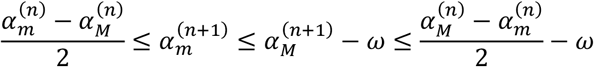

From 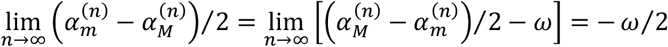,

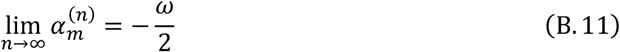

Accordingly,

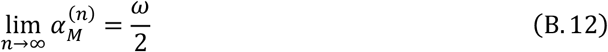

### Lemma 8

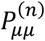 does not converge to zero as *n*→∞ for all *μ*.

**Proof:**

Lemma 8 is proved by contradiction.

Assumption 4: 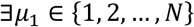 s.t. 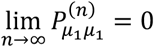

From Assumption 4 and Eq. (B. 4), 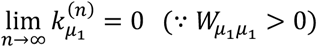.

Here, from Eq. (B. 12), 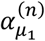 does not diverge to infinity as *n*→∞. Therefore, from the definition of 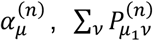 does not converge to zero. Accordingly, from 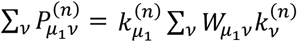 (∵Eq.(B.4)),

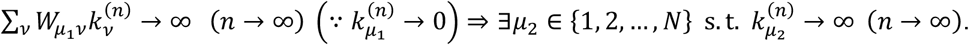

Then, from 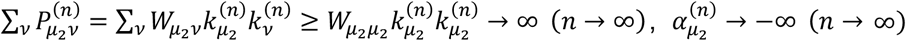. This is inconsistent with Eq. (B. 11). Therefore, Assumption 4 cannot be realized.

When 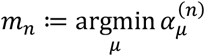 the followings are derived:

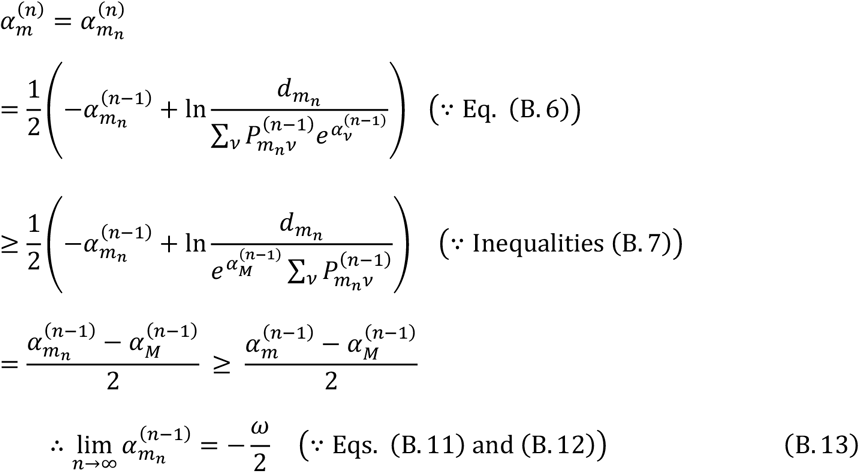

From 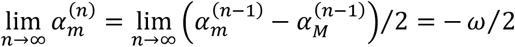 (∵ Eqs. (B. 11) and (B. 12)),

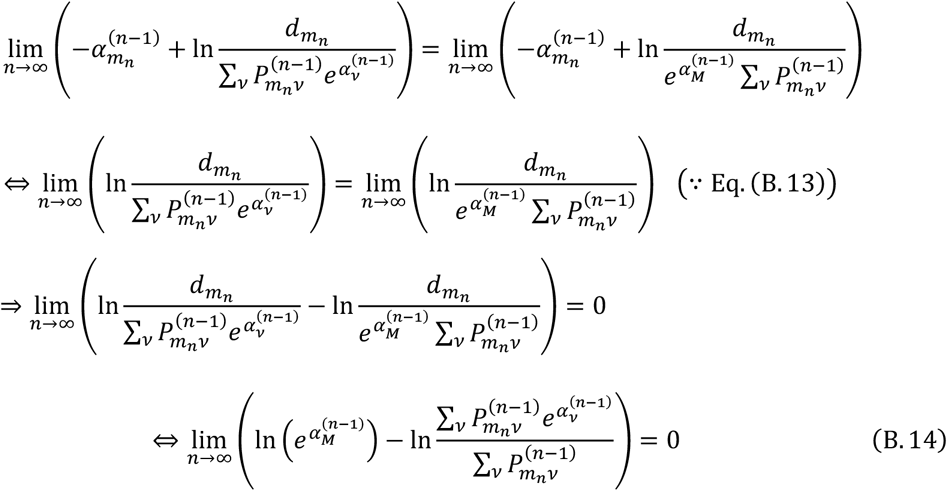

Here, 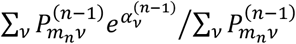 is a weighted average of 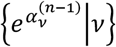. Because 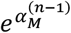 is the maximum value in 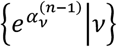 and 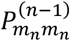 does not converge to zero as *n*→∞ (∵ Lemma 8), 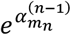 is necessary to approach 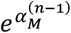 infinitely with *n*→∞ to satisfy Eq. (B. 14). Therefore, the following equations are valid:

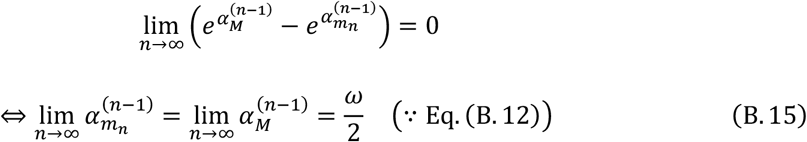

From Eqs. (B. 13) and (B. 15), *ω* = 0 (∵–*ω* = *ω*). Therefore, from Eqs. (B. 11) and (B. 12), Eq. (B. 5) is valid.

## Appendix C. Supplementary data

**Fig. C.1.**
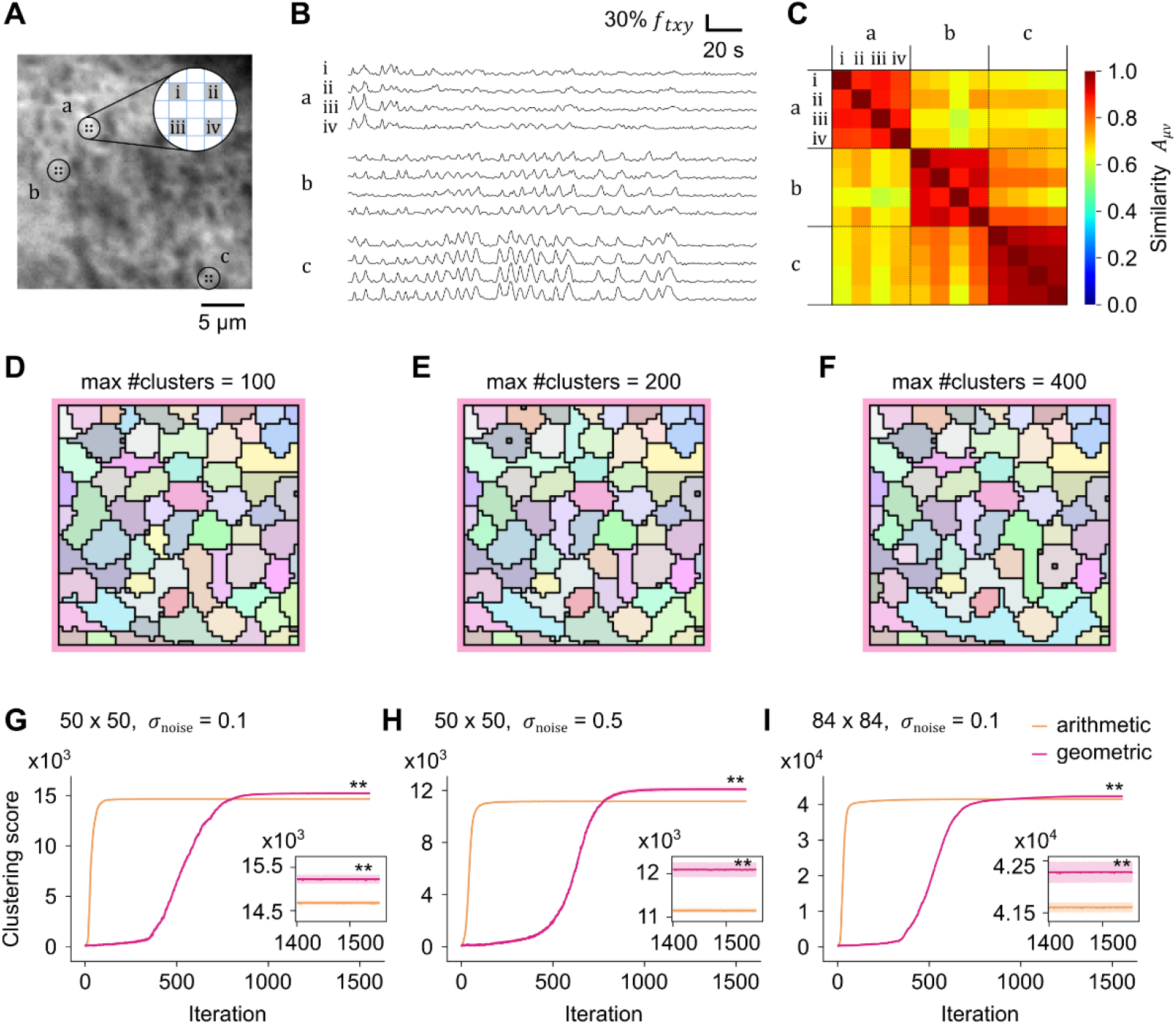
Validation of the cosine-based similarity and the geometric progression-based annealing schedule. (A) Basal fluorescence of the calcium imaging of the neuropil of a fly larva. Three boutons (a, b, and c) were marked. For each bouton, four pixels (i-iv) were sampled. (B) Preprocessed signals of the sampled pixels in (A). See section 2.2 for the preprocessing procedure. (C) Signal similarity between the pixels in (A), calculated as cosine between signal traces in (B), assumed as vectors. (D, E, and F) Clustering with a different maximum number of clusters (D: 100, E: 200, and F: 400). (G, H, and I) Clustering score development during simulated annealing. Magenta indicates geometric progression of inverse temperature beta, whereas orange indicates arithmetic progression. Beta reached 20 at the final iteration 1,557 in both progressions. Shaded regions show the standard deviation of five trials. The final clustering score of geometric progression was significantly larger than that of arithmetic progression (**: p < 10^-2^, Mann-Whitney U test). (G) Artificial data of 50 x 50 pixel array and sig_noise = 0.1. (H) Artificial data of 50 x 50 pixel array and *σ*_noise_ = 0.5. (I) Artificial data of 84 x 84 pixel array and *σ*_noise_ = 0.1.

**Fig. C.2.**
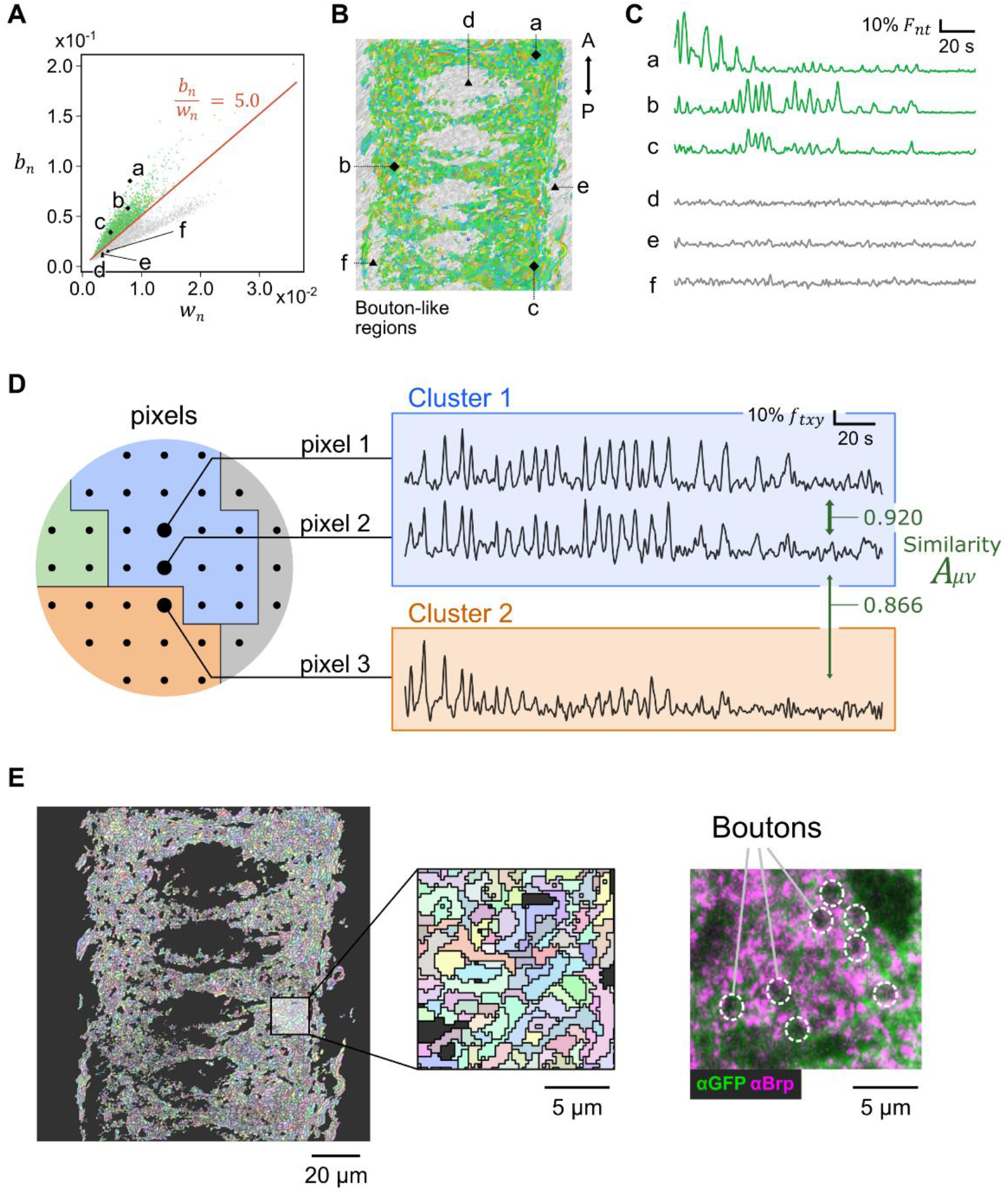
Pixels in imaging data were clustered into boutons. (A, B, and C) Detected bouton-like regions with sufficient (green) and insufficient (gray) S/N ratio from one sample. Regions a, b, and c showed sufficient S/N ratio, whereas d, e, and f didn’t. (A) Scatter diagram of bouton-like regions showing the intensity of noise (horizontal) and signals (vertical). Orange line is the threshold level of the S/N ratio. (B) Spatial configuration of the bouton-like regions. (C) Signal traces of the regions a-f. (D) Examples of activity traces of pixels. Left: Spatial configuration of pixels colored based on their cluster identity. Pixels 1 and 2 were included in cluster 1 (blue), whereas pixel 3 belonged to cluster 2 (orange). Right: Temporal traces of pixels 1, 2, and 3. The numbers colored in green are the values in the similarity matrix (see section 2.1.1.). (E) The sizes of extracted patches and actual boutons were comparable. Left: Bouton-like regions. Each region is shown by a different color. Middle: Expanded image of the white square in (E). Right: Immunostaining image of the neuropil with anti-GFP and anti-Brp (the marker of presynaptic sites), shown on the same scale as the middle panel. Dashed circles indicate boutons.

**Fig. C.3.**
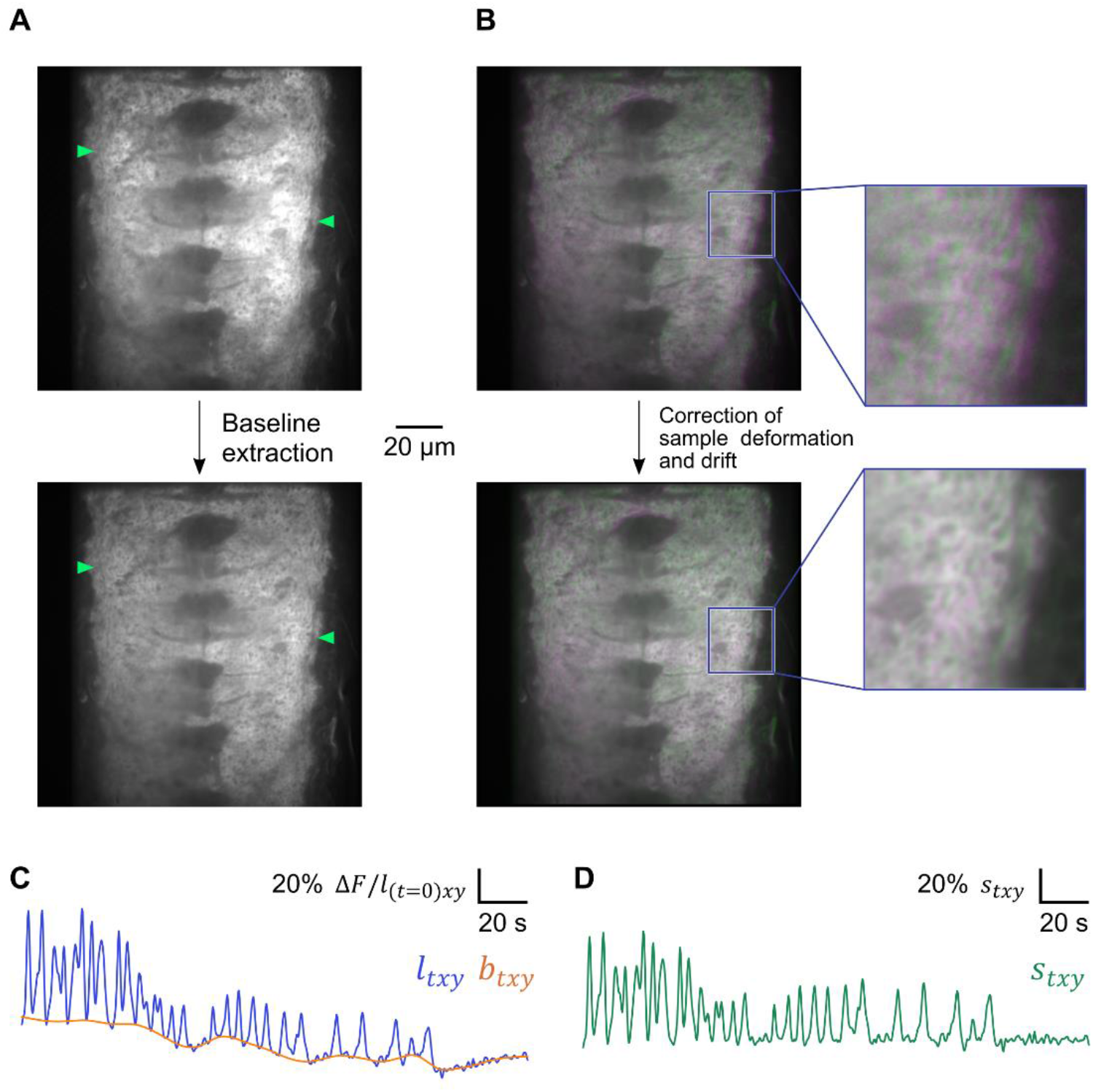
Baseline detection and correction. (A) An imaging frame showing neural activity (top) and the detected temporal baselines sectioned on the same frame (bottom). Top and bottom panels are shown in the same contrast. Green arrowheads indicate regions showing strong neural activity in the top panel. (B) Calcium imaging on two distinct frames (green: *t* = 245, magenta: *t* = 699) is merged. Top and bottom panels show images before and after correction of sample deformation and drift, respectively. Right panels are expansion images of left panels, showing misalignment was drastically reduced after the processing. (C) An activity trace of one pixel and detected baseline. The blue line is the original data, and the orange line is the detected baseline. (D) Baseline-corrected trace of the same pixel in (C).

**Fig. C.4.**
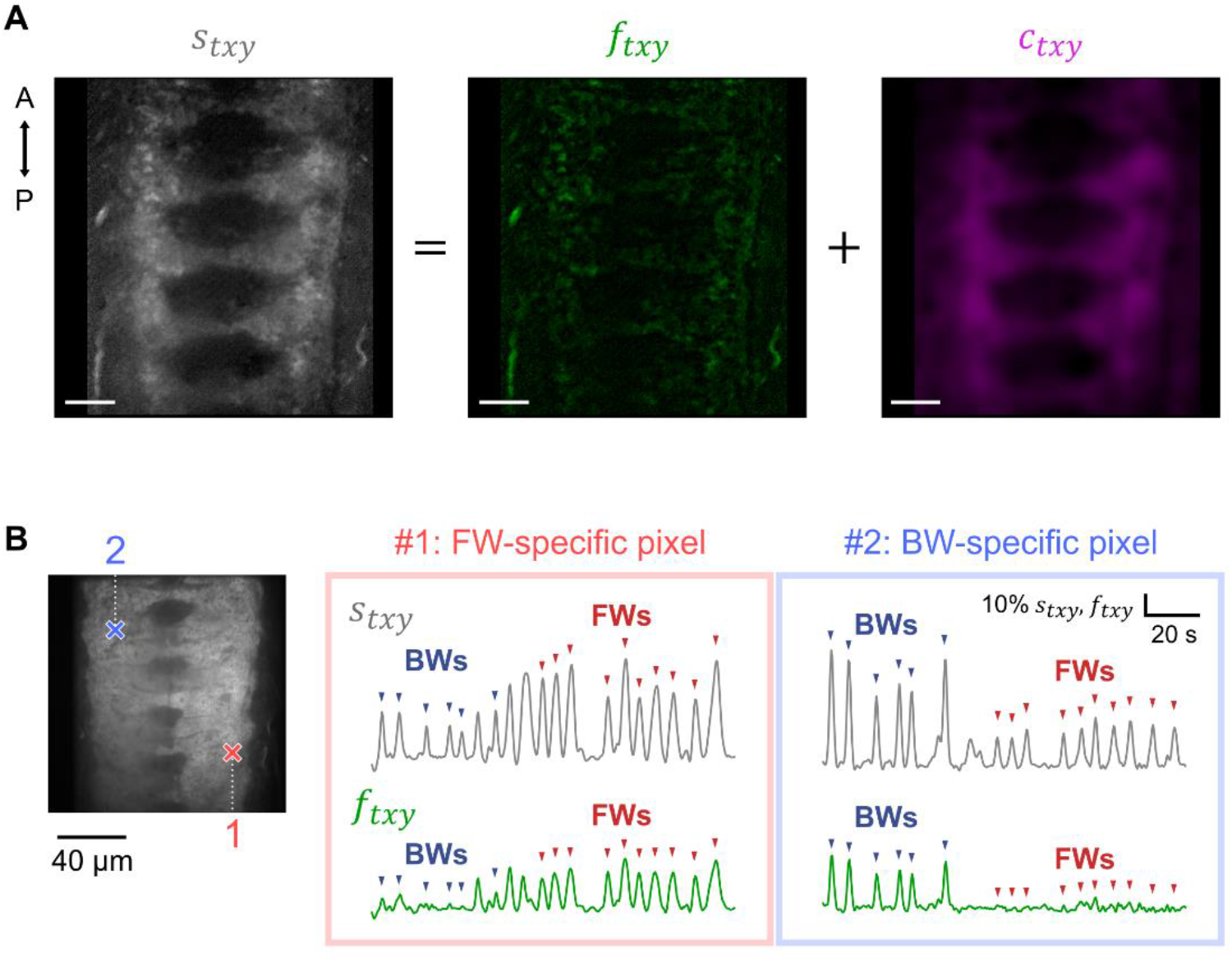
Calcium imaging data were successfully decomposed into well-focused and out-of-focus structures. (A) Fine-coarse decomposition on one example frame. Gray image shows original signals, while green and magenta images show decomposed fine and coarse components, respectively. Scale bar: 20 μm. (B) Forward- or backward-specific pixels can be recognized only in fine components. FW is a forward wave, and BW is a backward wave. Pixels #1 and #2 show FW-specific and BW-specific activity, respectively. Left: Spatial sites of pixels #1 and #2. Right: Temporal traces of pixels #1 and #2. Gray lines are original signals, while green lines are signals of extracted fine components. Red and blue arrowheads indicate FW and BW activity patterns, respectively.

**Fig. C.5.**
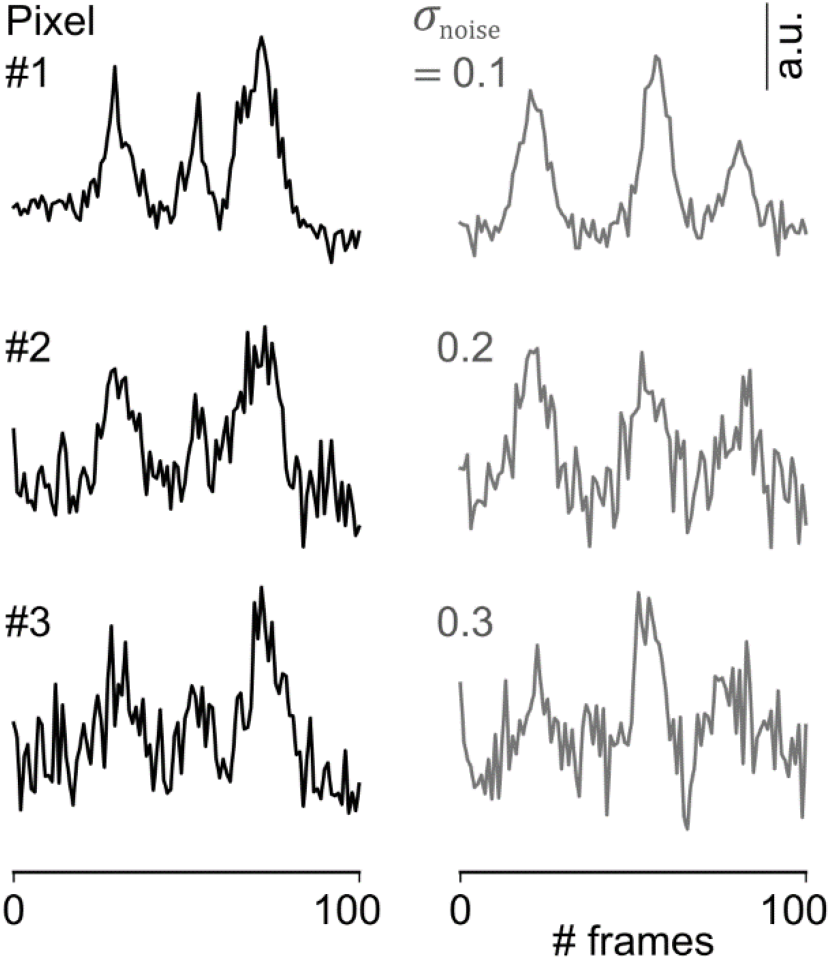
Comparison of biological calcium recording data and artificial data. Left: Temporal profiles of three example pixels in biological calcium imaging data. Right: Artificial burst-like signals of three distinct noise intensities (*σ*_noise_). The noise intensities of the real calcium imaging data corresponded to *σ*_noise_ = 0.1-0.3 in the artificial data.

## Notes

### Competing Interest Statement

The authors have declared no competing interest.

